# Structural Basis for Mis18 Complex Assembly: Implications for Centromere Maintenance

**DOI:** 10.1101/2021.11.08.466737

**Authors:** Reshma Thamkachy, Bethan Medina-Pritchard, Sang Ho Park, Carla G. Chiodi, Juan Zou, Maria de la Torre-Barranco, Kazuma Shimanaka, Maria Alba Abad, Cristina Gallego Páramo, Regina Feederle, Emilija Ruksenaite, Patrick Heun, Owen R. Davies, Juri Rappsilber, Dina Schneidman-Duhovny, Uhn-Soo Cho, A. Arockia Jeyaprakash

**Author notes:** Crystal structures Correspondence. These authors contributed equally (Reshma Thamkachy, Bethan Medina-Pritchard, Sang Ho Park).

## Abstract

The centromere, defined by the enrichment of CENP-A (a Histone H3 variant) containing nucleosomes, is a specialised chromosomal locus that acts as a microtubule attachment site. To preserve centromere identity, CENP-A levels must be maintained through active CENP-A loading during the cell cycle. A central player mediating this process is the Mis18 complex (Mis18α, Mis18β and Mis18BP1), which recruits the CENP-A specific chaperone HJURP to centromeres for CENP-A deposition. Here, using a multi-pronged approach, we characterise the structure of the Mis18 complex and show that multiple hetero- and homo-oligomeric interfaces facilitate the hetero-octameric Mis18 complex assembly composed of 4 Mis18α, 2 Mis18β and 2 Mis18BP1. Evaluation of structure-guided/separation-of-function mutants reveals structural determinants essential for Mis18 complex assembly and centromere maintenance. Our results provide new mechanistic insights on centromere maintenance, highlighting that while Mis18α can associate with centromeres and deposit CENP-A independently of Mis18β, the latter is indispensable for the optimal level of CENP-A loading required for preserving the centromere identity.

## Introduction

Faithful chromosome segregation during cell division requires bi-orientation of chromosomes on the mitotic spindle through the physical attachment of kinetochores to microtubules. Kinetochores are large multiprotein scaffolds that assemble on a special region of chromosomes known as the centromere (Musacchio and Desai, 2017, Cheeseman, 2014, Catania and Allshire, 2014, Fukagawa and Earnshaw, 2014). Whilst centromeres in some organisms, such as budding yeast, are defined by a specific DNA sequence, in most eukaryotes, centromeres are distinguished by an increased concentration of nucleosomes containing a histone H3 variant called CENP-A (Fukagawa and Earnshaw, 2014, McKinley and Cheeseman, 2016, Stellfox et al., 2013, Black et al., 2010). CENP-A containing nucleosomes recruit CENP-C and CENP-N, two proteins that are part of the constitutive centromere-associated network (CCAN) and that recruits the rest of the kinetochore components at the centromeric region of the chromosome (Carroll et al., 2010, Kato et al., 2013, Weir et al., 2016).

Whilst canonical histone loading is coupled with DNA replication, CENP-A loading is not (Dunleavy et al., 2011). This results in a situation where, after S-phase, the level of CENP-A nucleosomes at the centromere is halved due to the distribution of existing CENP-A to the duplicated DNA (Jansen et al., 2007, Dunleavy et al., 2009). To maintain centromere identity, centromeric CENP-A levels must be restored. This is achieved through active CENP-A loading at centromeres (during G1 in humans) via a pathway that requires the Mis18 complex (consisting of Mis18α, Mis18β and Mis18BP1) and the CENP-A chaperone, HJURP (Jansen et al., 2007, Fujita et al., 2007, Dunleavy et al., 2009, Foltz et al., 2009, Barnhart et al., 2011). The Mis18 complex can recognise and localise to the centromere, possibly through its proposed binding to CENP-C and/or other mechanisms which have not yet been identified (Dambacher et al., 2012, Stellfox et al., 2016, Moree et al., 2011). Once at the centromere, the Mis18 complex has been implicated in facilitating the deposition of CENP-A in several ways. There is evidence that the Mis18 complex affects DNA methylation and histone acetylation, which may facilitate CENP-A loading (Hayashi et al., 2004, Kim et al., 2012). But one of its most important and well-established roles of the Mis18 complex is the recruitment of HJURP, which binds a single CENP-A/H4 dimer and brings it to the centromere (Hu et al., 2011, Dunleavy et al., 2009, Barnhart et al., 2011). This then triggers a poorly understood process in which the H3 nucleosomes are removed and replaced with CENP-A nucleosomes. Finally, the new CENP-A nucleosomes are stably integrated into the genome, which requires several remodelled factors such as MgcRacGAP, RSF, Ect2, and Cdc42 (Lagana et al., 2010, Perpelescu et al., 2009).

The timing of CENP-A deposition is tightly regulated, both negatively and positively, by the kinases Cdk1 and Plk1, respectively, in a cell cycle-dependent manner (Silva et al., 2012, Spiller et al., 2017, Pan et al., 2017, McKinley and Cheeseman, 2014, Stankovic et al., 2017, Muller et al., 2014). Previous studies demonstrated that Cdk1 phosphorylation of Mis18BP1 prevents the Mis18 complex assembly and localisation to centromeres until the end of mitosis (when Cdk1 levels are reduced) (Spiller et al., 2017, Pan et al., 2017). Cdk1 also phosphorylates HJURP, which negatively regulates its binding to the Mis18 complex at the centromere (Muller et al., 2014, Stankovic et al., 2017, Wang et al., 2014). In cells, Plk1 is a positive regulator, and its activity is required for G1 centromere localisation of the Mis18 complex and HJURP. Plk1 has been shown to not only phosphorylate Mis18α/ý and Mis18BP1, but it has also been proposed to interact with phosphorylated Mis18 complex through its polo-box domain (PBD) (McKinley and Cheeseman, 2014).

As outlined above, a central event in the process of CENP-A deposition at centromeres is the Mis18 complex assembly. The Mis18 proteins, Mis18α and Mis18β, possess a well-conserved globular domain called the Yippee domain (also known as the MeDiY domain; spanning residues 77-180 in Mis18α and 73-176 in Mis18β) and C-terminal α-helices (residues 196-233 in Mis18α and 191-229 in Mis18β). We and others previously showed that the Yippee domains of Mis18 proteins can form a hetero-dimer, while the C-terminal helices form a hetero-trimer with two Mis18α and one Mis18β. However, the full-length proteins form a hetero-hexameric assembly with 4 Mis18α and 2 Mis18β. This led to a proposed model, where the Mis18α and Mis18β mainly interact via the C-terminal helices to form a hetero-trimer, and two such hetero-trimers interact via the Yippee hetero-dimerisation (Mis18α/Mis18β) or/and homo-dimerisation (Mis18α/Mis18α) to form a hetero-hexameric assembly (Nardi et al., 2016, Spiller et al., 2017, Pan et al., 2017, Pan et al., 2019).

Mis18BP1, the largest subunit of the Mis18 complex (1132 aa residues), is a multi-domain protein containing SANTA (residues 383-469) and SANT (residues 875-930) domains, which are known to have roles in regulating chromatin remodelling (Zhang et al., 2006, Aasland et al., 1996, Maddox et al., 2007). In-between these two domains resides the CENP-C binding domain (CBD) (Dambacher et al., 2012, Stellfox et al., 2016). *In vivo*, the CBD alone is not sufficient to recruit Mis18BP1 to the centromere and requires the N-terminus of the protein for proper localisation (Stellfox et al., 2016). We and others have previously shown that the N-terminal 130 amino acids of Mis18BP1 are sufficient for interaction with Mis18α/ý through their Yippee domains, and Cdk1 phosphorylation of Mis18BP1 at residues T40 and S110 inhibits its interaction with Mis18α/ý to form an octamer complex consisting of 2 Mis18BP1, 4 Mis18α and 2 Mis18β (Spiller et al., 2017, Pan et al., 2017).

Although the importance of the Mis18 complex assembly and function is well appreciated, structural understanding of the intermolecular interfaces responsible for the Mis18 complex assembly and their functions are yet to be identified. Here, we have characterised the structural basis of the Mis18 complex assembly using an integrative structure modelling approach that combines X-ray crystallography, Electron Microscopy (EM), Small Angle X-ray Scattering (SAXS), Cross-Linking Mass Spectrometry (CLMS), AlphaFold and computational modelling. By evaluating the structure-guided mutations *in vitro* and *in vivo*, we provide important insights into the key structural elements responsible for Mis18 complex assembly and centromere maintenance.

## Results

### Structural basis for the assembly of Mis18α/ý core modules

Mis18α and Mis18β possess two distinct but conserved structural entities, a Yippee domain and a C-terminal α-helix (**Fig. 1a** **and S1a & b**). Mis18α possesses an additional α-helical domain upstream of the Yippee domain (residues 39-76). Previous studies have shown that Mis18α Yippee domain can form a homo-dimer or a hetero-dimer with Mis18β Yippee domain whereas Mis18α/ý C-terminal helices form a robust 2:1 heterotrimer (Subramanian et al., 2016, Spiller et al., 2017, Pan et al., 2017). Disrupting Yippee homo- or hetero-dimerisation in full-length proteins, while did not abolish their ability to form a complex, did perturb the dimerisation of Mis18α/ý heterotrimer (Spiller et al., 2017). Contrarily, intermolecular interactions involving the C-terminal helices of Mis18α and Mis18β are essential for Mis18α/ý complex assembly (Nardi et al., 2016). Overall, the available biochemical data suggest the presence of at least three independent structural core modules within the Mis18α/ý complex: the Mis18α Yippee homo-dimer, the Mis18α/ý Yippee hetero-dimer and the Mis18α/ý C-terminal helical assembly. Here, we structurally characterised these modules individually and together as a holo-complex.

**Figure 1:**
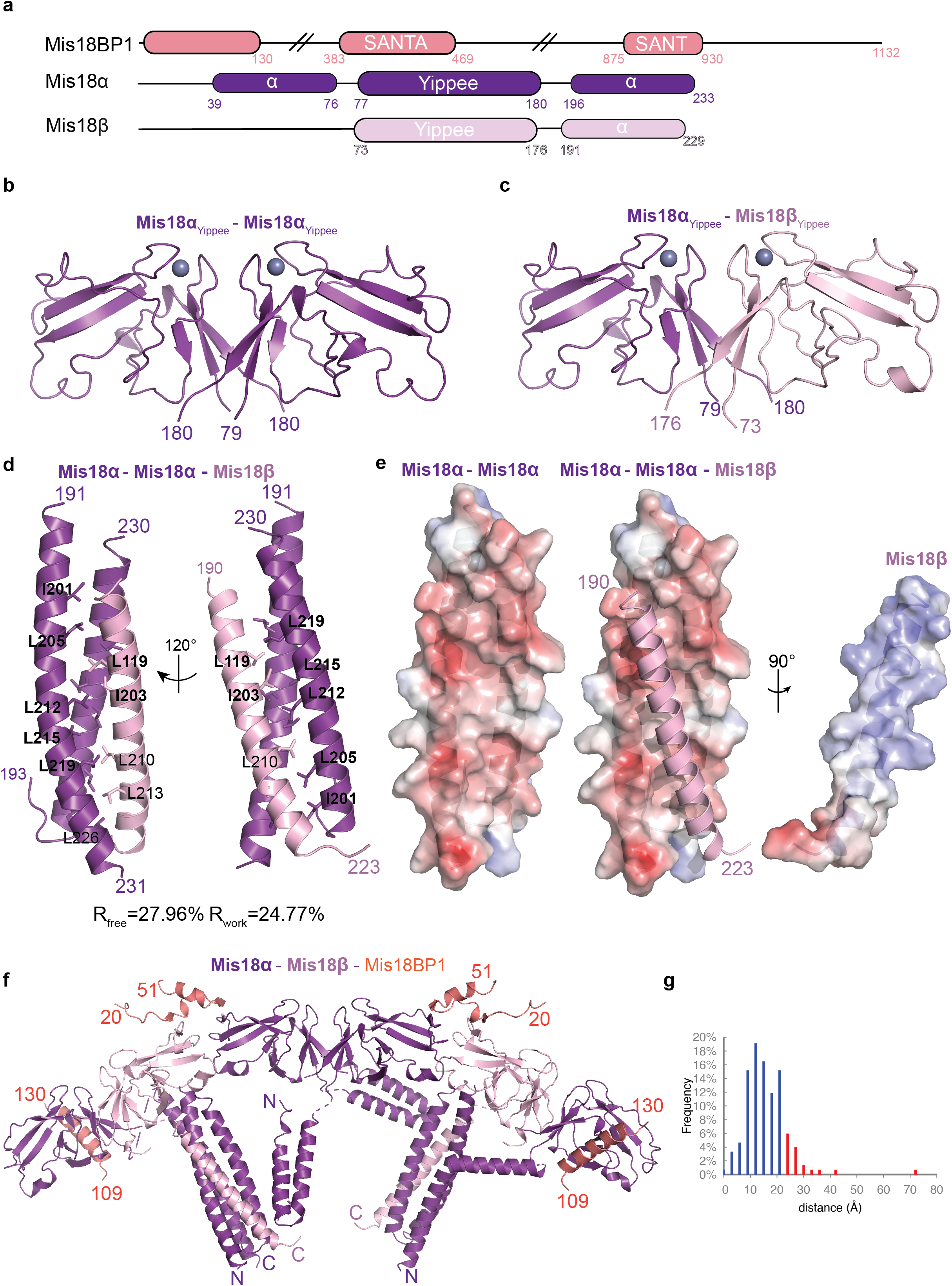
Mis18α/ý Contains Two Independent Structural Domains that can Oligomerise. **a)** Schematic representation of structural features of Mis18BP1 (salmon), Mis18α (purple) and Mis18ý (light pink). Filled boxes represent folded domains. SANTA and SANT domain boundaries as defined in UniProt (Q6P0N0). **b)** Cartoon representation of the crystal structure of human Mis18α_Yippee_ homo-dimer (PDB ID: 7SFZ). **c)** Cartoon representation of the human Mis18α_Yippee_/Mis18ý_Yippee_ hetero-dimer modelled by homology to the structure in Fig. 1b. Mis18α is shown in purple and Mis18ý in light pink (modelled using Phyre2, www.sbg.bio.ic.ac.uk/phyre2/ (Kelley et al., 2015)). **d)** Cartoon representation of the crystal structure of Mis18α_C-term_/Mis18ý_C-term_ (PDB ID: 7SFY). Mis18α is shown in purple and Mis18ý in light pink. **e)** Mis18α_C-term_ domains are shown in surface representation and coloured based on electrostatic surface potential calculated using APBS (Baker et al., 2001). Mis18ý_C-term_ shown as cartoon. **f)** Model of the Mis18_core_ complex generated using partial structures determined using X-ray crystallography and AlphaFold2 (Jumper et al., 2021) and cross-linking restrained molecular docking in EM maps. Mis18BP1 shown in salmon, Mis18α in purple and Mis18ý in light pink. **g)** Histograms show the percentage of satisfied or violated cross-links for structures modelled using MODELLER (Sali and Blundell, 1993).

#### Mis18α Yippee homo-dimer

We previously determined a crystal structure of the Yippee domain in the only homologue of Mis18 in *S. pombe* (PDB: 5HJ0), showing that it forms a homo-dimer (Subramanian et al., 2016). To determine the structure of human Mis18 Yippee domains, we purified and crystalised Mis18α_Yippee_ (residues 77-190). The crystals diffracted X-rays to about 3 Å resolution, and the structure was determined using the molecular replacement method. The final model was refined to R and R_free_ factors of 20.26% and 25.00%, respectively (**Table S1** and **Fig. 1b**, PDB ID: 7SFZ). The overall fold of the Mis18α_Yippee_ is remarkably similar to the previously determined *S. pombe* Mis18_Yippee_ homo-dimer structure (Subramanian et al., 2016). In brief, the monomeric Mis18α_Yippee_ is formed by two antiparallel β-sheets that are held together by a Zn^2+^ ion coordinated via loops containing C-X-X-C motifs. The Mis18α_Yippee_ dimerisation is mediated via a back-to-back arrangement of a ‘three-stranded’ β-sheet from each monomer.

#### Mis18α/ý Yippee hetero-dimer

As repeated efforts to crystallise the Mis18α/ý Yippee heterodimer were not successful, using the Mis18α_Yippee_ as a template we generated high-confidence structural models for the Mis18α/ý_Yippee_ hetero-dimer using Raptorx (http://raptorx.uchicago.edu/) (Källberg et al., 2012) and AlphaFold2 (Jumper et al., 2021) (**Fig. 1c**). As observed for Mis18α_Yippee_ homo-dimer, the Mis18α/ý Yippee hetero-dimerisation is also mediated via the back-to-back arrangement of the three-stranded beta sheets of Mis18α and Mis18β Yippee domains.

#### Mis18α/ý C-terminal helical assembly

Previous studies have shown that recombinantly purified C-terminal α-helices of Mis18α and Mis18β form a hetero-trimer with 2 copies of Mis18α and 1 copy of Mis18β (Spiller et al., 2017, Pan et al., 2017). However, in the absence of high-resolution structural information, how Mis18 C-terminal helices interact to form a hetero-trimer and how the structural arrangements of α-helices influence the relative orientations of the Yippee domains, and hence the overall architecture of the Mis18α/ý hexamer assembly, remained unclear. We purified Mis18α spanning aa residues 191 to 233 and Mis18β spanning aa residues 188 and 229 (**Fig. 1a, S1a & b**) and crystallised the reconstituted complex. The crystals diffracted X-rays to about 2.5 Å resolution. The structure was determined using single wavelength anomalous dispersion method. After iterative cycles of refinement and model building, the final model was refined to R and R_free_ factors of 24.77% and 27.96%, respectively (**Table S1,** PDB ID: 7SFY). The asymmetric unit contained two copies of Mis18α/ý hetero-trimer. The final model included Mis18α residues 191 to 231 in one copy, Mis18α residues 193 to 230 in the second copy, and Mis18β residues 190 to 223 (**Fig. 1d**). The two Mis18α helices interact in an antiparallel orientation, and one helix is stabilised in a slightly curved conformation. This arrangement results in a predominantly negatively charged groove that runs diagonally on the surface formed by the Mis18α helices (**Fig. 1d** **& e**). This observation is consistent with the theoretically calculated pI of the Mis18α helix (pI=4.9). In contrast, the pI of the Mis18β helix is 8.32. This charge complementarity appears to facilitate the interaction with Mis18α, as a positively charged surface of the Mis18β helix snugly fits in the negatively charged groove of the Mis18α/α interface. A closer look at the intermolecular interactions reveals tight hydrophobic interactions along the ‘spine’ of the binding groove with electrostatic interactions ‘zipping-up’ both sides of the Mis18β helix (**Fig 1e**). The binding free energy calculated based on the buried accessible surface area suggests a nanomolar affinity interaction between the helices of Mis18α and Mis18β. It should be noted that the crystal structure presented here differs from the previously predicted models in terms of either the subunit stoichiometry (Pan et al., 2019) or the directional arrangement of individual subunits (Mis18α and Mis18β in parallel orientation with the 2^nd^ Mis18α in an anti-parallel orientation (this work) vs all parallel (Pan et al., 2019)).

### Multiple surfaces of Mis18α/ý Yippee hetero-dimers contribute to the overall oligomeric assembly of the Mis18 complex

Full-length Mis18α/ý complex or the Mis18_core_ complex (Mis18α - Mis18β - Mis18BP1_20-130_) were not amenable for structural characterisation using X-ray crystallography possibly due to their intrinsic flexibility. Consistent with this notion, the SAXS profiles collected for the Mis18α/ý ýN (Mis18α residues 77-187 and Mis18β residues 56-183), Mis18α/ý and Mis18_core_ complexes suggest that these complexes possess an elongated shape with flexible features (**Fig. S2, Table S2**). Hence, to understand the overall assembly of the Mis18 complex we took an integrative structure modelling approach, combining the crystal structures of Mis18α_Yippee_ dimer and Mis18α/Mis18β C-terminal hetero-trimeric helical assembly together with the homology/AlphaFold modelling of Mis18α_Yippee_/Mis18ý_Yippee_ hetero-dimer, negative staining EM, SAXS and CLMS analysis of the Mis18_core_ complex.

The negative staining electron micrographs of the Mis18_core_ complex cross-linked using GraFix (Kastner et al., 2008) revealed a good distribution of particles (**Fig. S3a**). Particle picking, followed by a few rounds of 2D classifications revealed classes with defined structural features (**Fig. S3b**). Some of the 2D projections resembled the shape of a ‘handset’ of a telephone with bulkier ‘ear’ and ‘mouth’ pieces. Differences in the relative orientation of bulkier features of the 2D projection suggested conformational heterogeneity. The three-dimensional volumes calculated for the particles were similar (approximately 220 x 105 x 80 Å) and in agreement with the *D_max_* calculated from SAXS analysis (**Fig. S2d**).

We attempted to assemble the whole Mis18 complex using AlphaFold-multimer (AFM), with full length Mis18α (in purple), Mis18β (in pink) and two small region of Mis18BP1 (20-51 and 109-130; in salmon) (Evans et al., 2021). The AFM converged towards a structure with six Yippee domains stacked in a line-like arrangement in the Mis18α_Yippee_-Mis18ý_Yippee_-Mis18α_Yippee_-Mis18α_Yippee_-Mis18ý_Yippee_-Mis18α_Yippee_ order and two triple helix bundles, each formed by C-terminal α-helices of 2 copies of Mis18β and 1 copy of Mis18β. However, the modelled two helical bundles had all three helices in a parallel orientation that is not supported by our crystal structures (**Fig. 1d**) and crosslinks (**Fig. S2e**). We modified the relative orientation of the helices to match the crystal structure by superposing the latter on the AFM model (**Fig. 1f, 1g & S3d**). Using crosslinks and docking we have added the N-terminal helices of the Mis18α. Cross-linking data indicates that these helices have multiple orientations with respect to the rest of the structure, contacting both Yippee domains and triple helix bundles. The linker between the Yippee domain and the C-terminal helix is the shortest in Mis18β (**Fig. 1a**), further supporting the arrangement of the Yippee domains within the assembly. The integrative model of the Mis18 complex fits well in the EM map. Interestingly, the serial arrangement of the Yippee domains utilises the second Yippee dimerisation interface observed in the crystal packing of both human Mis18α Yippee and *S. pombe* Mis18 Yippee (**Fig. S3d, highlighted by zoom in view**). Accordingly, disrupting this interface by mutating Mis18α residues C154 and D160 (**Fig. S3d**) perturbed Mis18 oligomerisation as evidenced by SEC analysis (**Fig**. **S3e**).

### Mis18α oligomerisation via the C-terminal helical bundle assembly is essential for Mis18α/ý centromere localisation and new CENP-A loading

Although the subunit stoichiometry and the arrangement of Mis18α/ý C-terminal helices within the helical bundle proposed by Nardi et al. 2016 are different from the data presented here, the Mis18α residues (I201, L205, L212, L215 and L219) that were predicted by them to stabilise the helical bundle do indeed form the ‘spine’ of the hydrophobic core running along the triple helical bundle (**Fig. 1d** and **e**). Mutating these residues perturbed the ability of Mis18α tethered at an ectopic LacO site to facilitate CENP-A deposition at the tethering site (Nardi et al., 2016). However, how these Mis18α mutants perturb the oligomeric structure of the Mis18α/ý C-terminal helical bundle and how this structural perturbation affects CENP-A loading at endogenous centromeres remain as open questions.

To address these questions, we first tested these mutants using *in vitro* amylose pull-down assays by mixing recombinantly purified WT and mutant His-MBP-Mis18β_188-229_ and His-SUMO-Mis18α_191-233_ proteins. Mutating these residues to Ala (Mis18α_I201A/L205A_ and Mis18α_L212A/L215A/L219A_) or Asp (Mis18α_I201D/L205D_) abolished the ability of Mis18α α-helix to interact with Mis18β_188-229_ (**Fig. S4a**). SEC MALS analysis of His-SUMO tagged Mis18α_188-233_ showed that on its own, Mis18α WT protein can form a dimer, whilst introducing I201A/L205A or L212A/L215A/L219A results in both proteins forming a monomer (**Fig. S4c**). Co-immunoprecipitation (Co-IP) assays using an anti-Mis18α antibody were performed on cells where endogenous Mis18α was depleted, and Mis18α−mCherry was co-expressed with Mis18β-GFP to check for complex formation (**Fig. S4b**). In line with our *in vitro* pull-downs, the co-IPs using a Mis18α antibody revealed that Mis18α_WT_-mCherry interacted with Mis18β-GFP while Mis18α_I201A/L205A_ and Mis18α_L212A/L215A/L219A_ mutants did not (**Fig. S4b**). To evaluate the role of this interaction on centromere localisation of Mis18α and Mis18β and CENP-A deposition, these mutants were further tested in HeLa cells.

HeLa CENP-A-SNAP cells (McKinley and Cheeseman, 2014) were depleted of endogenous Mis18α by siRNA (**Fig. S4d**) and simultaneously rescued with either WT or mutant Mis18α-mCherry (**Fig. S4e**), then visualised by immunofluorescence along with ACA. Unlike Mis18α_WT_, the Mis18α mutants (Mis18α_I201A/L205A_, Mis18α_I201D/L205D_ and Mis18α_L212A/L215A/L219A_) all failed to localise to centromeres (**Fig. 2a**). As expected, Mis18β-GFP co-expression showed co-localisation between Mis18ý_WT_ with Mis18α_WT_. However, in cells expressing Mis18α_I20A1/L205A_, Mis18α_I201D/L205D_ and Mis18α_L212A/L215A/L219A_, Mis18β could no longer co-localise with Mis18α at the centromere. Together, this confirms that Mis18β depends on its interaction with Mis18α and the formation of the C-terminal triple helical assembly to localise at centromeres.

**Figure 2:**
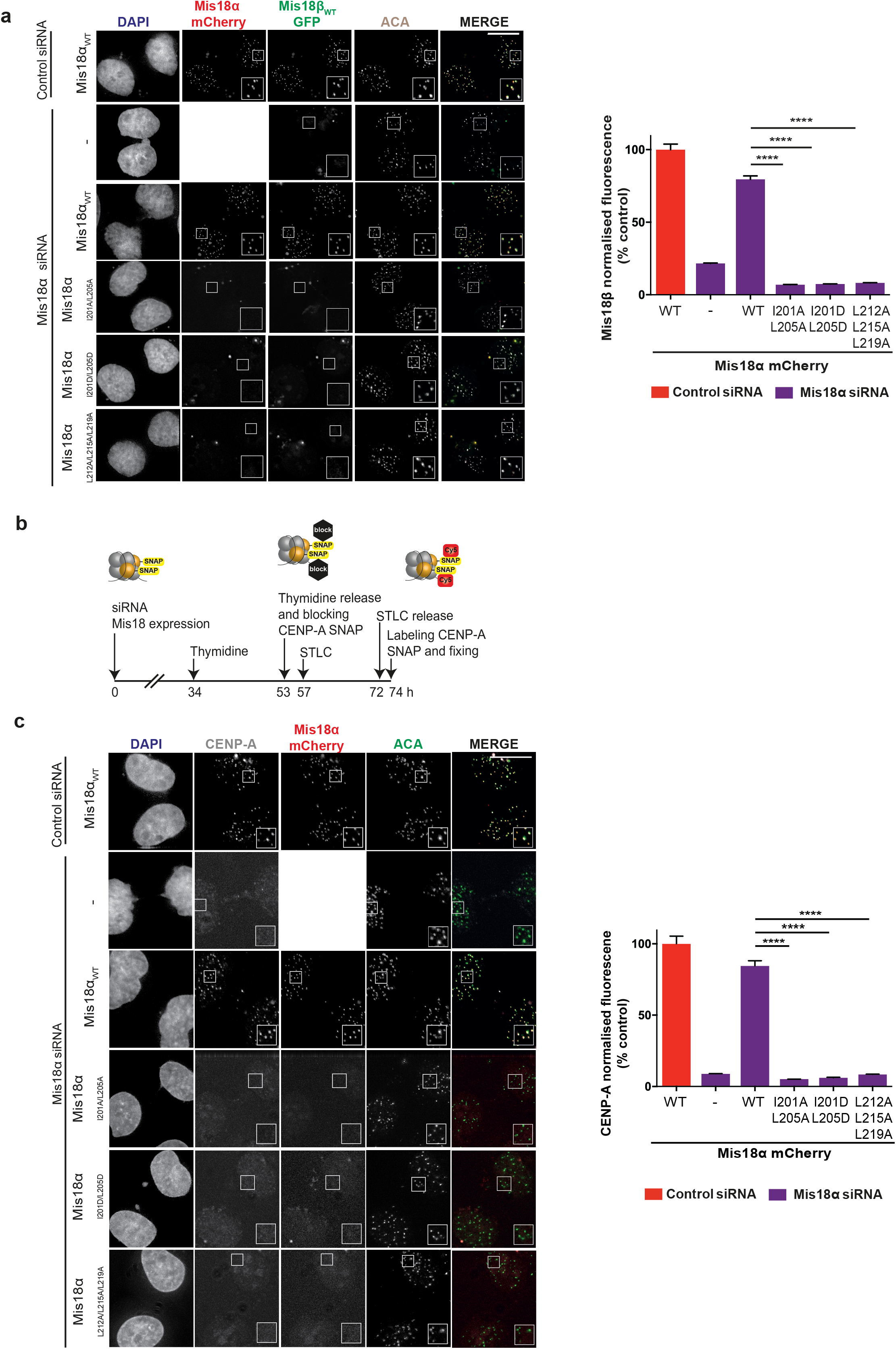
Mis18α Mutations Disrupting the Mis18α/ý Triple Helical Assembly Result in Loss of Mis18α/ý Centromere Localisation and CENP-A Deposition. **a)** Representative fluorescence images (left panel) and quantification (right panel) assessing the ability of Mis18α_WT_-mCherry, Mis18α_I201A/L205A_-mCherry, Mis18α_I201D/L205D_-mCherry and Mis18α_L212A/L215A/L219A-_mCherry to co-localise with Mis18ý GFP at endogenous centromeres in HeLa (Mann-Whitney U test; \*\*\*\**P* < 0.0001, *n* ≥ 1236). Cells were co-transfected with either control or Mis18α siRNA, as stated, in 3 independent experiments. Error bars show ±SEM. Scale bars, 10 μm. All conditions have been normalised to control conditions: cells transfected with control siRNA and Mis18α_WT_-mCherry. **b)** Schematic representation of the experimental set-up used to evaluate the effect of Mis18α and Mis18ý mutants on new CENP-A-SNAP loading. **c)** Representative fluorescence images (left panel) and quantification (right panel) assessing the ability of Mis18α_WT_-mCherry, Mis18α_I201A/L205A_-mCherry, Mis18α_I201D/L205D_-mCherry and Mis18α_L212A/L215A/L219A_-mCherry to deposit new CENP-A-SNAP at endogenous centromeres (Mann-Whitney U test; \*\*\*\**P* < 0.0001, *n* ≥ 886). Cells were co-transfected with either control or Mis18α siRNA, as stated, in 3 independent experiments. Error bars show ±SEM. Scale bars, 10 μm. All conditions have been normalised to control conditions: cells transfected with control siRNA and Mis18α_WT_-mCherry.

We then evaluated the impact of Mis18α mutants not capable of forming the C-terminal helical bundle on new CENP-A deposition. We did this by performing a Quench-Chase-Pulse CENP-A-SNAP Assay according to Jansen *et al*. (Jansen et al., 2007) (**Fig. 2c**). HeLa CENP-A-SNAP cells were depleted of endogenous Mis18α and rescued with either Mis18α_WT_ or Mis18α mutants (Mis18α_I20A1/L205A_, Mis18α_I201D/L205D_ and Mis18α_L212A/L215A/L219A_). The existing CENP-A was blocked with a non-fluorescent substrate of the SNAP, and the new CENP-A deposition in the early G1 phase was visualised by staining with the fluorescent substrate of the SNAP. Mis18α_WT_ rescued new CENP-A deposition to levels compared to that of control siRNA (**Fig. 2b and c**). However, Mis18α_I20A1/L205A_, Mis18α_I201D/L205D_ and Mis18α_L212A/L215A/L219A_ abolished new CENP-A loading almost completely, indicating that the formation of the Mis18 triple helical bundle is essential for CENP-A deposition (**Fig. 2c**).

### Mis18α associates with the centromere independently of Mis18β and can deposit CENP-A, but efficient CENP-A loading requires Mis18

We again performed amylose *in vitro* pull-down assays, using His-SUMO-Mis18α_191-233 WT_ and mutant His-MBP-Mis18β_188-229_ proteins, to assess the ability of Mis18β mutant to form a triple-helical bundle with Mis18α. Based on our X-ray crystal structure (**Fig. 1d**), we identified one cluster (L199/I203) in Mis18β and observed that mutating these residues to either Ala (Mis18ý_L199A/I203A_) or Asp (Mis18ý_L199D/I203D_) either reduced or abolished its ability to interact with Mis18α_191-233_ (**Fig. 3a**). Co-IP analysis using an anti-Mis18α antibody was performed on cells where endogenous Mis18β was depleted, and Mis18β-GFP was expressed along Mis18α-mCherry to check for complex formation. Western blot analysis showed that Mis18ý_WT_ could interact with Mis18α-mCherry and that the ability of Mis18ý_L199D/I203D_ to interact with Mis18α was reduced (**Fig. 3a**, right panel).

**Figure 3:**
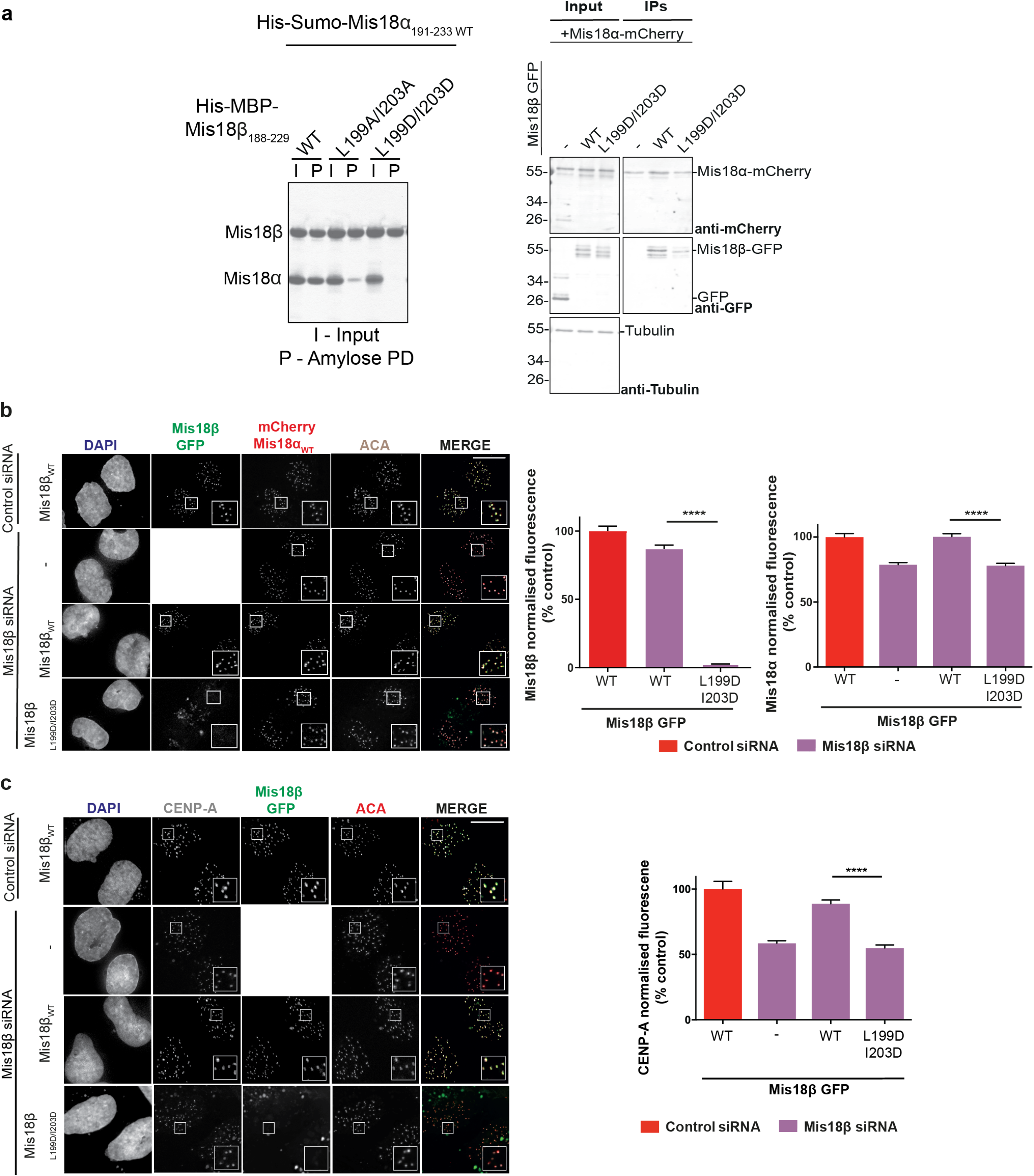
Mis18α Associates with Centromeres in a Mis18ý-Independent Manner but Requires Mis18ý for Efficient CENP-A Loading. **a)** Left panel shows SDS-PAGE analysis of cobalt and amylose pull-down of His-MBP-Mis18ý_188–229 WT_ and mutants with His-SUMO-Mis18α_191–233_. SDS-PAGE shows protein bound to nickel resin as input (I) and protein-bound to amylose resin to assess interaction (P). Right panel shows Western blot analysis of co-immunoprecipitation (Co-IP) experiments using Mis18α antibody to test interaction of Mis18α−mCherry and Mis18ý GFP with and without mutations in the C-terminal α-helices. Top panel shows blot against mCherry, middle panel shows blot against GFP and bottom panel shows blot against tubulin as loading control. **b)** Representative fluorescence images (left panel) and quantification (right panel) used to evaluate the ability of Mis18ý_WT_-GFP and Mis18ý_L199D/I203D_-GFP to co-localise with mCherry-Mis18α at endogenous centromeres. Middle panel, quantification of Mis18ý signal. Right panel, quantification of Mis18α signal (Mann-Whitney U test; \*\*\*\**P* < 0.0001, *n* ≥ 927). **c)** Representative fluorescence images (left panel) and quantification (right panel) used to evaluate the ability of Mis18ý_WT_-GFP and Mis18ý_L199D/I203D_ GFP to deposit new CENP-A-SNAP at endogenous centromeres (Mann-Whitney U test; \*\*\*\**P* < 0.0001, *n* ≥ 947). Cells were co-transfected with either control or Mis18ý siRNA, as stated, in 3 independent experiments. Error bars show ±SEM. Scale bars, 10 μm. All conditions have been normalised to control conditions: cells transfected with control siRNA and Mis18ý_WT_-GFP.

To assess the contribution of Mis18β for the centromere association and function of Mis18α, we evaluated the Mis18β mutant (Mis18ý_L199D/I203D_), which cannot form the triple helical assembly with Mis18α, in siRNA rescue assays by expressing Mis18β-GFP tagged proteins in a mCherry-Mis18α cell line (McKinley and Cheeseman, 2014). Depletion of endogenous Mis18β and simultaneous transient expression of Mis18ý_WT_-GFP led to co-localisation of Mis18β with Mis18α at centromeres (**Fig. 3b, S4d & S4e**). Under these conditions, Mis18ý_WT_-GFP levels at centromeres were comparable to that of the control siRNA. Whereas Mis18ý_L199D/I203D_ failed to localise at the centromeres. Strikingly, Mis18ý_L199D/I203D_ perturbed centromere association of Mis18α only moderately (**Fig 3b**,). This suggests that Mis18α can associate with centromeres in a Mis18β independent manner.

Next, we assessed the contribution of Mis18β for CENP-A deposition in the Quench-Chase-Pulse CENP-A-SNAP assay described above. Endogenous Mis18β was depleted using siRNA, and Mis18ý_WT_ and Mis18ý_L199D/I203D_ were transiently expressed as GFP-tagged proteins in HeLa cells expressing CENP-A-SNAP. Mis18ý_WT_ rescued new CENP-A deposition to comparable levels to the ones observed in the control siRNA-Mis18β WT condition (**Fig 3c**). Interestingly, unlike the Mis18α mutants (Mis18α_I20A1/L205A_, Mis18α_I201D/L205D_ and Mis18α_L212A/L215A/L219A_), Mis18ý_L199D/I203D_ did not abolish new CENP-A loading but reduced the levels only moderately.

Together, these analyses demonstrate that Mis18α can associate with centromeres and deposit new CENP-A independently of Mis18β. However, efficient CENP-A loading requires Mis18β.

### Structural basis for centromere recruitment of Mis18α/ý by Mis18BP1

Previous studies have shown that Mis18BP1 N-terminus (1-130 aa) is required to bind Mis18α/ý (Spiller et al., 2017). However, how Mis18α/ý Yippee domains recognise Mis18BP1 is not clear. Our structural analysis suggests that two Mis18BP1 fragments, a short helical segment spanning aa residues 110-130 (Mis18BP1_110-130_) and a region spanning aa residues 24-50 (Mis18BP1_24-50_) interact with Mis18α Yippee domain and with an interface formed between Mis18α/ý Yippee hetero-dimers, respectively (**Fig. 4a**). Mis18BP1_110-130_ binds at a hydrophobic pocket of the Mis18α Yippee domain formed by amino acids L83, F85, W100, I110, V172 and I175. This hydrophobic pocket is surrounded by hydrophilic amino acids E103, D104, T105, S169 E171 facilitating additional electrostatic interactions with Mis18BP1_110-130_

**Figure 4:**
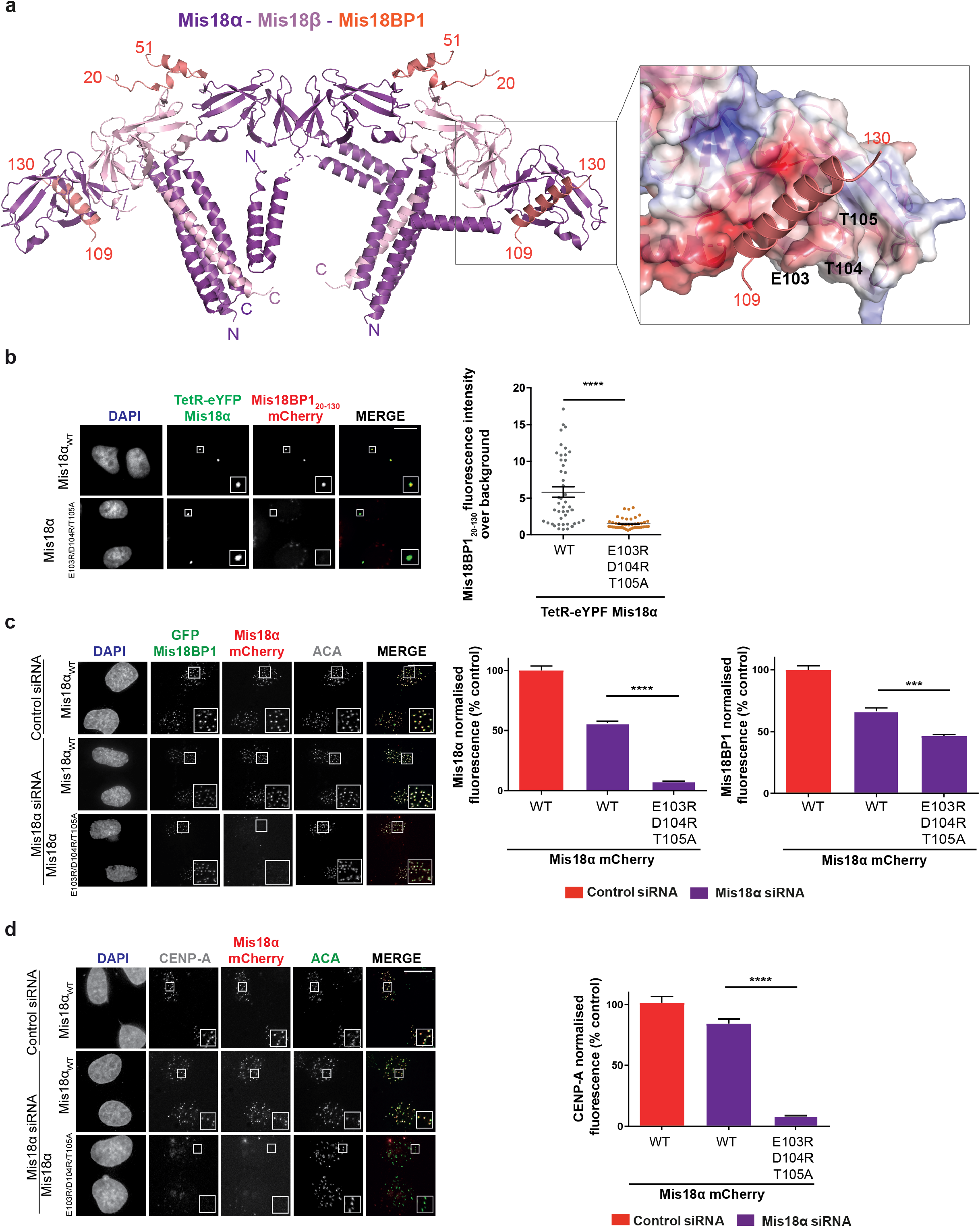
Disrupting the Mis18BP1 Binding Interface of Mis18α Prevents its Centromere Localisation and CENP-A Deposition. **a)** Mis18α/Mis18β model and its surface representation coloured based on electrostatic surface potential (zoom panel), highlighting the residues proposed to be involved in Mis18BP1 binding. Mis18α shown in purple, Mis18β shown in light pink and Mis18BP1 shown in salmon. **b)** Representative images and quantification showing the recruitment of either Mis18BP1_20-130_-mCherry by different Mis18α constructs (WT and mutant) tethered to the alphoid^tetO^ array in HeLa 3-8. Tethering of TetR-eYFP-Mis18α_WT_ and TetR-eYFP-Mis18α_E103R/D104R/T105A_ testing recruitment of Mis18BP1_20-130_ mCherry (Mann-Whitney U test; \*\*\*\**P* < 0.0001, *n* ≥ 45). Date from 3 independent experiments. Central lines show mean whilst error bars show SEM. Scale bars, 10 μm. **c)** Representative fluorescence images (left panel) and quantifications (right panel) evaluating the ability of Mis18α_WT_-mCherry and Mis18α_E103R/D104R/T105A_ to co-localise with GFP-Mis18BP1 at endogenous centromeres. Middle panel, quantification of Mis18α signal and right panel, quantification of Mis18BP1 signal (Mann-Whitney U test; \*\*\*\**P* < 0.0001, *n* ≥ 856). **d)** Representative fluorescence images (left panel) and quantifications (right panel) evaluating the ability of Mis18α_WT_-mCherry and Mis18α_E103R/D104R/T105A_ to deposit new CENP-A-SNAP at endogenous centromeres (Mann-Whitney U test; \*\*\**P* = 0.0001, \*\*\*\**P* < 0.0001, *n* ≥ 896). Cells were co-transfected with either control or Mis18α siRNA, as stated, in 3 independent experiments. Error bars show ±SEM. Scale bars, 10 μm. All conditions have been normalised to control conditions: cells transfected with control siRNA and Mis18α_WT_-mCherry.

(**Fig. 4a**). Mis18BP1_24-50_ contains two short β strands that interact at Mis18α/ý Yippee interface extending the six-stranded-β sheets of both Mis18α and Mis18β Yippee domains. Notably, the two Cdk1 phosphorylation sites on Mis18BP1 (T40 and S110) that we and others have shown to disrupt Mis18 complex assembly (Spiller et al., 2017, Pan et al., 2017) lie directly within the Mis18 α/ý binding interface predicted by this model, providing the structural basis for Cdk1 mediated regulation of Mis18 complex assembly. Consistent with this model, several cross-links observed between Mis18BP1 and Mis18α and Mis18β map to these residues. Mutating the negatively charged amino acid cluster of Mis18α (E103, D104 and T105) that is juxtaposed to Mis18BP1_110-130_ in a TetR-eYFP-Mis18α vector (TetR-eYFP-Mis18α_E103R/D104R/T105R_) transfected in HeLa cells with an ectopic synthetic alphoid^tetO^ array integrated in a chromosome arm significantly perturbed Mis18α’s ability to recruit Mis18BP1_20-130_-mCherry to the tethering site as compared to Mis18α_WT_ (**Fig. 4b**).

Furthermore, we probed the effects of perturbing Mis18α-Mis18BP1 interaction on endogenous centromeres. We depleted Mis18α in a cell line that stably expresses CENP-A-SNAP and allows inducible expression of GFP-Mis18BP1 (McKinley and Cheeseman, 2014). We then assessed the ability of transfected Mis18α-mCherry to co-localise with Mis18BP1 at centromeres. Depletion of Mis18α and simultaneous expression of either Mis18α_WT_-mCherry or Mis18α_E103R/D104R/T105A_-mCherry revealed that, unlike Mis18α_WT,_ Mis18α_E103R/D104R/T105A_ failed to localise at endogenous centromeres (**Fig. 4c**, middle panel). We also observed a slight decrease in the levels of GFP-Mis18BP1 at the centromere when Mis18α_E103R/D104R/T105A_ was expressed as compared to Mis18α_WT_ (**Fig. 4c**, right panel). Consistent with the observation of reduced centromeric Mis18α, when Mis18α_E103R/D104R/T105A_-mCherry is expressed, the quantification of new CENP-A deposition in HeLa cell expressing CENP-A-SNAP showed a significant reduction of new CENP-A deposition at the centromere indicating that the interaction of Mis18α with Mis18BP1 is essential for centromeric recruitment of the Mis18 complex and for CENP-A loading (**Fig. 4d**).

## Discussion

Mis18 complex assembly is a central process essential for the recruitment of CENP-A/H4 bound HJURP and the subsequent CENP-A deposition at centromeres (Jansen et al., 2007, Fujita et al., 2007, Dunleavy et al., 2009). Thus far, several studies, predominantly biochemical and cellular, have characterised interactions and functions mediated by the two distinct structural domains of the Mis18 proteins, the Yippee and C-terminal α-helical domains of Mis18α and Mis18β (Spiller et al., 2017, Pan et al., 2017, Nardi et al., 2016, Stellfox et al., 2016). Some of the key conclusions of these studies include: (1) Mis18α/ý is a hetero-hexamer made of 4 Mis18α and 2 Mis18β; (2) The Yippee domains and C-terminal α-helices of Mis18α and Mis18β have the intrinsic ability to homo- or hetero-oligomerise, and form three distinct oligomeric modules in different copy numbers – a Mis18α_Yippee_ homo-dimer, two copies of Mis18α/ý_Yippee_ hetero-dimers and two hetero-trimers made of Mis18α/ý C-terminal helices (2 Mis18α and 1 Mis18β); (3) the two copies of Mis18α/ý_Yippee_ hetero-dimers each bind one Mis18BP1_20-130_ and form a hetero-octameric Mis18_core_ complex (Mis18α/Mis18β/Mis18BP1_20-130_: a Mis18α/ý hetero-hexamer bound to 2 copies of Mis18BP1_20-130_). However, no experimentally determined structural information is available for the human Mis18 complex. This is crucial to identify the amino acid residues essential for the assembly of Mis18α/ý and the holo-Mis18 complexes and to determine the specific interactions that are essential for the localisation of Mis18 complex to centromeres and its function.

Here, we have taken an integrative structural approach that combines X-ray crystallography, electron microscopy and homology modelling with cross-linking mass spectrometry to characterise the structure of the Mis18 complex. Our analysis shows that Mis18α/ý hetero-trimer is stabilised by the formation of a triple helical bundle with a Mis18α/ý_Yippee_ hetero-dimer on one end and Mis18α_Yippee_ monomer on the other. Two such Mis18α/ý hetero-trimers assemble as a hetero-hexamer via the homo-dimerisation of the Mis18α_Yippee_ domains. The crystal structure of Mis18α/ý_C-term_ triple helical structure allowed us to design several separation of function Mis18α and Mis18β mutants. These mutations specifically perturb the ability of Mis18α or Mis18β to assemble into the helical bundle, while retaining their other functions, if there are any. Functional evaluation of these mutants in cells has provided important new insights into the molecular interdependencies of the Mis18 complex subunits. Particularly, the observations that: (1) Mis18α can associate with centromeres and deposit CENP-A independently of Mis18β, and (2) depletion of Mis18β or disrupting the incorporation of Mis18β into the Mis18 complex, while does not abolish CENP-A loading, reduces the CENP-A deposition amounts, questions the consensus view that Mis18α and Mis18β always function as a single structural entity to exert their function to maintain centromere maintenance.

Whilst proteins involved in CENP-A loading have been well established, the mechanism by which the correct levels of CENP-A are controlled is yet to be thoroughly explored and characterised. The data presented here suggest that Mis18β mainly contributes to the quantitative control of centromere maintenance – by ensuring the right amounts of CENP-A deposition at centromeres – and maybe one of several proteins that control CENP-A levels. Future studies will focus on dissecting the mechanisms underlying the Mis18β-mediated control of CENP-A loading amounts along with any other mechanisms involved.

Previous studies using siRNA to deplete Mis18α shows that is does not effect Mis18BP1 localisation and that Mis18BP1 can associate with centromeres independently of Mis18α (McKinley and Cheeseman, 2014). The separation of function Mis18α mutant unable to bind Mis18PB1, characterised here, shows that disrupting Mis18α-Mis18BP1 interaction completely abolishes Mis18α’s ability to associate with centromeres and new CENP-A loading. This highlights that Mis18BP1-mediated centromere targeting is the major centromere recruitment pathway for the Mis18α/ý complex.

Previously published work identified amino acid sequence similarity between the N-terminal region of Mis18α and R1 and R2 repeats of the HJURP that mediates Mis18α/ý interaction (Pan et al., 2019). Deletion of the Mis18α N-terminal region enhanced HJURP interaction with the Mis18 complex. This led to speculation that the N-terminal region of Mis18α might directly interact with the HJURP binding site of the Mis18 complex and thereby modulating HJURP binding. Our work presented here strengthens this speculation and provides the structural justification. We show that the N-terminal helical region of Mis18α makes extensive contacts with the C-terminal helices of Mis18α and Mis18β that mediate HJURP binding. In the future, it will be important to address how and when the interference caused by the N-terminal region of Mis18α is relieved for efficient HJURP binding by the Mis18 complex.

## Acknowledgements

We would like to thank David Kelly from the Centre Optical Instrumentation Laboratory, Martin Wear from the Edinburgh Protein Production Facility as well as Marcus Wilson and Maarten Tuijtel of the Cryo-Electron Microscopy Facility for their help. We also thank Diamond Light Source and the staff of beamline B21 (proposal sm23510), as well as Advanced Photon Source and the staff at the beamlines LS-CAT 21 ID-G and ID-D. Thanks also to Iain Cheeseman for the kind gift of cell lines. The Wellcome Trust generously supported this work through Senior Research Fellowships to A.A. Jeyaprakash (202811), J. Rappsilber (084229), O. Davies (219413/Z/19/Z) and P. Heun (103897/Z/14/Z), a Centre Core Grant (092076 and 203149) and an instrument grant (108504) to the Wellcome Trust Centre for Cell Biology. AAJ and his team are co-funded by the European Union (ERC Advanced Grant, CHROMSEG, 101054950). Views and opinions expressed are, however, those of the authors only and do not necessarily reflect those of the European Union or the European Research Council.

Neither the European Union nor the granting authority can be held responsible for them. The work of D. Schneidman is supported by ISF 1466/18 and Israeli Ministry of Science and Technology.

## Author Contributions

A.A.J. and U.S.C. conceived the project. R.T., B.M.P., S.H.P., M.A.A and A.A.J. designed the experiments. S.H.P. and K.S. preformed crystal structure characterisation. C.G.C. and M.T.B. preformed EM characterisation. B.M.P. and J.Z. preformed cross-linking analysis. C.G.P. and O.R.D. analysed SAXS data. D.S.D. and A.A.J. preformed modelling. R.T. and S.H.P. preformed biochemical characterisation. R.T. and B.M.P. preformed *in vivo* assays and analysis. R.F. and E.R. generated and validated anti-bodies. J.R. and P.H. provided resources, expertise, and feedback. B.M.P., C.G.C., R.T., D.S.D. and A.A.J. wrote the manuscript.

The authors declare no competing financial interests.

## Material and Methods

### Plasmids

For crystallisation, a polycistronic expression vector for the C-terminal coiled-coil domains of Mis18α (residues 191-233, Mis18α_C-term_) and Mis18β (residues 188-229, Mis18ý_C-term_) were produced with the N-terminal 6His-SUMO-(His-SUMO) and 6His-MBP-tags (His-MBP), respectively. Mis18α_Yippee_ (residues 77-190) was cloned into the pET3a vector with the N-terminal 6His-tag.

For all other recombinant proteins, codon optimised sequences (GeneArt) for Mis18α and Mis18β were cloned into pET His6 TEV or pET His6 msfGFP TEV (9B Addgene plasmid #48284, 9GFP Addgene plasmid #48287, a kind gift from Scott Gradia), respectively. They were combined to make a single polycistronic plasmid. The boundaries of ΔN for Mis18α and Mis18β were 77-187 and 56-183. Mis18BP1_20-130_ was cloned in pEC-K-3C-His-GST and pET His6 MBP TEV (9C Addgene plasmid #48286).

Non-codon optimised sequences were amplified from a human cDNA library (MegaMan human transcription library, Agilent). Mis18α, Mis18β and Mis18BP1_20-130_ were cloned into pcDNA3 mCherry LIC vector, pcDNA3 GFP LIC vector (6B Addgene plasmid #30125, 6D Addgene plasmid #30127, a kind gift from Scott Gradia) and TetR-eYFP-IRES-Puro vector as stated. All mutations were generated following QuikChange site-directed mutagenesis protocol (Stratagene).

### Expression and purification of recombinant proteins

For crystallisation, both Mis18α/ý_C-term_ domains and Mis18α_Yippee_ were transformed and expressed in *Escherichia coli* BL21 (DE3) using the auto-inducible expression system (Studier, 2005). The cells were harvested and resuspended in the lysis buffer containing 30 mM Tris-HCl pH7.5, 500 mM NaCl, and 5 mM β-mercaptoethanol with protease inhibitor cocktails. The resuspended cells were lysed using the ultra-sonication method and centrifuged at 20,000 x g for 50 min at 4°C to remove the cell debris. After 0.45 μm filtration of the supernatant, the lysate was loaded into the cobalt affinity column (New England Biolabs) and eluted with a buffer containing 30 mM Tris-HCl pH7.5, 500 mM NaCl, 5 mM ý-mercaptoethanol, and 300 mM imidazole. The eluate was loaded into the amylose affinity column (New England Biolabs) and washed with a buffer containing 30 mM Tris-HCl pH7.5, 500 mM NaCl, and 5 mM β-mercaptoethanol. To cleave the His-MBP tag, on-column cleavage was performed by adding Tobacco Etch Virus (TEV) protease (1:100 ratio) into the resuspended amylose resin and incubated overnight at 4°C. The TEV cleavage released the untagged Mis18α/ý_C-term_ domains in solution, and the flow through fraction was collected and concentrated using a Centricon (Millipore). The protein was loaded onto a HiLoad™ 16/600 Superdex™ 200 column (GE Healthcare) equilibrated with a buffer containing 30 mM Tris-HCl pH7.5, 100 mM NaCl, and 1 mM TCEP. To further remove the contaminated MBP tag, the sample was re-applied into the amylose affinity column, and the flow-through fraction was collected and concentrated to 20 mg/ml for the crystallisation trial. SeMet (selenomethionine) incorporated Mis18α/ý_C-term_ domains were expressed with PASM-5052 auto-inducible media (Studier, 2005). The SeMet-substituted Mis18α/ý_C-term_ domains were purified using the same procedure described above.

The purification of His tagged Mis18α_Yippee_ employed the same purification method used for Mis18α/ý_C-term_ domains except for the amylose affinity chromatography step. The purified Mis18α_Yippee_ from the HiLoad™ 16/600 Superdex™ 200 chromatography was concentrated to 13.7 mg/ml with the buffer containing 30 mM Tris-HCl pH7.5, 100 mM NaCl, and 1 mM TCEP.

All other proteins were expressed in *Escherichia coli* BL21 (*DE3*) Gold cells using LB. After reaching an O.D. ∼ 0.6 at 37°C, cultures were cooled to 18°C and induced with 0.35 mM IPTG overnight. The His-Mis18α/His-GFP-Mis18β complex was purified by resuspending the pellet in a lysis buffer containing 20 mM Tris-HCl pH 8.0 at 4°C, 250 mM NaCl, 35 mM imidazole pH 8.0 and 2 mM β-mercaptoethanol supplemented with 10 μg/ml DNase, 1mM PMSF and cOmplete™ EDTA-free (Sigma). After sonication, clarified lysates were applied to a 5 ml HisTrap™ HP column (GE Healthcare) and washed with lysis buffer followed by a buffer containing 20 mM Tris-HCl pH 8.0 at 4°C, 1 M NaCl, 35 mM imidazole pH 8.0, 50 mM KCl, 10 mM MgCl_2_, 2 mM ATP and 2 mM β-mercaptoethanol and then finally washed with lysis buffer. The complex was then eluted with 20 mM Tris-HCl pH 8.0 at 4°C, 250 mM NaCl, 500 mM imidazole pH 8.0 and 2 mM β-mercaptoethanol. Fractions containing proteins were pooled, and TEV was added (if needed) whilst performing overnight dialyses against 20 mM Tris-HCl pH 8.0 at 4°C, 150 mM NaCl and 2 mM DTT.

His-GST-Mis18BP1_20-130_ was purified in the same manner as above with the following modifications: the lysis and elution buffers contained 500 mM NaCl, whilst the dialysis buffer contained 75 mM NaCl. His-MBP-Mis18BP1_20-130_ was purified using the same lysis buffer containing 500 mM NaCl and purified using amylose resin (NEB). Proteins were then eluted by an elution buffer containing 10 mM Maltose.

If needed, proteins were subjected to anion exchange chromatography using the HiTrap™ Q column (GE Healthcare) using the ÄKTA™ start system (GE Healthcare). Concentrated fractions were then injected onto either Superdex™ 75 increase 10/300 or Superdex™ 200 increase 10/300 columns equilibrated with 20 mM Tris-HCl pH 8.0 at 4°C, 100-250 mM NaCl and 2 mM DTT using the ÄKTA™ Pure 25 system (GE Healthcare).

### Interaction trials

Pull-down assays used to test the interaction between the C-terminus of Mis18α and Mis1β were performed by initially purifying the proteins through the cobalt affinity chromatography, as described for wild type proteins, and the eluted fractions were loaded into the amylose affinity resin, pre-equilibrated with a binding buffer consisting of 30 mM Tris-HCl pH7.5, 500 mM NaCl, and 5 mM β-mercaptoethanol. Amylose resins were washed with the binding buffer, and the proteins were eluted with a binding buffer containing 20 mM maltose. The fractions were subjected to SDS-PAGE analysis.

Pull-down assay using the amylose resin to test interactions between Mis18α/μ and Mis18BP1_20-130_ were done as described previously (Pan et al., 2017). Briefly, purified proteins were diluted to 10 μM in 40 μl binding buffer, 50 mM HEPES pH 7.5, 1 M NaCl, 1 mM TCEP, 0.01% Tween® 20. One third of the mixture was taken as input, and the remaining fraction was incubated with 40 μl amylose resin for 1 h at 4°C. The bound protein was separated by washing with binding buffer three times, and the input and bound fractions were analysed by SDS-PAGE.

### Crystallisation, data collection, and structure determination

Purified Mis18α/μ_C-term_ domains and Mis18α_Yippee_ were screened and crystallised using the hanging-drop vapour diffusion method at room temperature with a mixture of 0.2 μl of the protein and 0.2 μl of crystallisation screening solutions. The crystals of Mis18α/μ_C-term_ domains were grown within a week with a solution containing 0.2 M magnesium acetate and 20% (w/v) PEG 3350. SeMet-substituted Mis18α/μ_C-term_ domains crystals were grown by the micro-seeding method with a solution containing 0.025 M magnesium acetate and 14% (w/v) PEG 3350. The crystals of SeMet-substituted Mis18α/μ_C-term_ domains were further optimised by mixing 1 μl of the protein and 1 μl of the optimised crystallisation solution containing 0.15 M magnesium acetate and 20% (w/v) PEG 3350. The crystals of Mis18α_Yippee_ were obtained in 2 M ammonium sulfate, 2% (w/v) PEG 400, and 100 mM HEPES at pH 7.5. The crystals of Mis18α/m_C-term_ domains and Mis18α_Yippee_ were cryoprotected with the crystallisation solutions containing 20% and 25% glycerol, respectively. The cryoprotected crystals were flash-frozen in liquid nitrogen. Diffraction datasets were collected at the beamline LS-CAT 21 ID-G and ID-D of Advanced Photon Source (Chicago, USA). The data set were processed and scaled using the DIALS (Winter et al., 2018) via Xia2 (Winter et al., 2013). The initial model of Mis18α/ý_C-term_ domains was obtained using the SAD method with SeMet-derived data using the Autosol program (Terwilliger, 2000). The molecular replacement of the initial model as a search model against native diffraction data was performed using the Phaser program within the PHENIX program suite (Liebschner et al., 2019). The initial model of Mis18α_Yippee_ was calculated by molecular replacement method (Phaser) using yeast Mis18 Yippee-like domain structure (PDB ID: 5HJ0) (Subramanian et al., 2016) as a search model. The final structures were manually fitted using the Coot program (Emsley and Cowtan, 2004) and the refinement was carried out using REFMAC5 (Afonine et al., 2010). The quality of the final structures was validated with the MolProbity program (Chen et al., 2010).

### SEC-MALS

Size-exclusion chromatography (ÄKTA-MicroTM, GE Healthcare) coupled to UV, static light scattering and refractive index detection (Viscotek SEC-MALS 20 and Viscotek RI Detector VE3580; Malvern Instruments) was used to determine the molecular mass of protein and protein complexes in solution. Injections of 100 µl of 2–6 mg/ml material were used. His-SUMO-Mis18α_188-233_ (∂A_280nm_/∂c = 0.43 AU.ml.mg^-1^) WT and mutants were run on a Superdex 75 increase 10/300 GL size exclusion column pre-equilibrated in 50 mM HEPES pH 8.0, 150 mM NaCl and 1 mM TCEP at 22°C with a flow rate of 1.0 ml/min.

Light scattering, refractive index (RI) and A_280nm_ were analysed by a homo-polymer model (OmniSEC software, v5.02; Malvern Instruments) using the parameters stated for the protein, ∂n/∂c = 0.185 ml.g^-1^ and buffer RI value of 1.335. The mean standard error in the mass accuracy determined for a range of protein-protein complexes spanning the mass range of 6-600 kDa is ± 1.9%.

### SAXS

SEC-SAXS experiments were performed at beamline B21 of the Diamond Light Source synchrotron facility (Oxfordshire, UK). Protein samples at concentrations >5 mg/ml were loaded onto a Superdex™ 200 Increase 10/300 GL size exclusion chromatography column (GE Healthcare) in 20 mM Tris pH 8.0, 150 mM KCl at 0.5 ml/min using an Agilent 1200 HPLC system. The column outlet was fed into the experimental cell, and SAXS data were recorded at 12.4 keV, detector distance 4.014 m, in 3.0 s frames. Data were subtracted, averaged and analysed for Guinier region *Rg* and cross-sectional *Rg* (*Rc*) using ScÅtter 3.0 (http://www.bioisis.net), and *P(r)* distributions were fitted using *PRIMUS* (Konarev et al., 2003). *Ab-initio* modelling was performed using *DAMMIN* (Svergun, 1999), in which 30 independent runs were performed in P1 or P2 symmetry and averaged.

### Gradient fixation (GraFix)

Fractions from the gel filtration peak were concentrated to 1 mg/mL using a Vivaspin® Turbo (Sartorius) centrifugal filter, and the buffer exchanged into 20 mM HEPES pH8.0, 150 mM NaCl, and 2 mM DTT for GraFix (Kastner et al., 2008, Stark, 2010). A gradient was formed with buffers A, 20 mM HEPES pH 8.0, 150 mM NaCl, 2 mM DTT, and 5% sucrose and B, 20 mM HEPES pH 8.0, 150 mM NaCl, 2 mM DTT, 25% sucrose, and 0.1% glutaraldehyde using the Gradient Master (BioComp Instruments). 500 μl of sample was applied on top of the gradient, and the tubes centrifuged at 40,000 rpm at 4°C using a Beckman SW40 rotor for 16 h. The gradient was fractionated in 500 μl fractions from top to bottom, and the fractions were analysed by SDS-PAGE with Coomassie blue staining and negative staining EM.

### Negative staining sample preparation, data collection and processing

Copper grids, 300 mesh, with continuous carbon layer (TAAB) were glow-discharged using the PELCO easiGlow™ system (Ted Pella). GraFix fractions with and without dialysis were used. Dialysed fractions were diluted to 0.02 mg/ml. 4 μl of sample were adsorbed for 2 min

onto the carbon side of the glow-discharged grids, then the excess was side blotted with filter paper. The grids were washed in two 15 μl drops of buffer and one 15 μl drop of 2% uranyl acetate, blotting the excess between each drop, and then incubated with a 15 μl drop of 2% uranyl acetate for 2 min. The excess was blotted by capillary action using a filter paper, as previously described (Scarff et al., 2018).

The grids were loaded into a Tecnai F20 (Thermo Fisher Scientific) electron microscope, operated at 200 kV, field emission gun (FEG), with pixel size of 1.48 Å. Micrographs were recorded using an 8k x 8k CMOS F816 camera (TVIPS) at a defocus range of -0.8 to -2μm. For Mis18α/ý/Mis18BP1_20-130_ (Mis18_core_), 163 micrographs were recorded and analysed using CryoSPARC 3.1.0 (Punjani et al., 2017). The contrast transfer function (CTF) was estimated using Gctf (Zhang, 2016). Approximately 750 particles were manually picked and submitted to 2D classification. The class averages served as templates for automated particle picking. Several rounds of 2D classification were employed to remove bad particles and assess the data, reducing the 14,840 particles to 5,540. These were used to generate three *ab-initio* models followed by homogeneous refinement with the respective particle sets.

### CLMS

Cross-linking was performed on gel filtered complexes dialysed into PBS. 16 μg EDC and 35.2 μg sulpho-NHS were used to cross-link 10 μg of Mis18α/ý with Mis18BP1_20-130_ (Mis18_core_) for 1.5 h at RT. The reactions were quenched with final concentration 100 mM Tris–HCl before separation on Bolt™ 4–12% Bis-Tris Plus gels (Invitrogen). Sulfo-SDA (sulfosuccinimidyl 4,4’-azipentanoate) (Thermo Scientific Pierce) cross-linking reaction was a two-step process. First, sulfo-SDA mixed with Mis18α/ý (0.39 μg/μl) at different ratio (w/w) of 1:0.07, 1:0.13, 1:0.19, 1:0.38, 1:0.5, 1:0.75, 1:1 and 1:1.4 (Mis18α/ý:Sulfo-SDA) was allowed to incubate 30 min at room temperature to initiate incomplete lysine reaction with the sulfo-NHS ester component of the cross-linker. The diazirine group was then photoactivated for 20 mins using UV irradiation from a UVP CL-1000 UV Cross-linker (UVP Inc.) at 365 nm (40 W). The reactions were quenched with 2 μl of 2.7 M ammonium bicarbonate before loading on Bolt™ 4–12% Bis-Tris Plus gels (Invitrogen) for separation. Following previously established protocol [38], either the whole sample or specific bands were excised, and proteins were digested with 13 ng/μl trypsin (Pierce) overnight at 37°C after being reduced and alkylated. The digested peptides were loaded onto C18-Stage-tips [39] for LC-MS/MS analysis.

LC-MS/MS analysis was performed using an Orbitrap Fusion Lumos (Thermo Fisher Scientific) coupled on-line with an Ultimate 3000 RSLCnano system (Thermo Fisher Scientific) with a “high/high” acquisition strategy. The peptide separation was carried out on a 50cm EASY-Spray column (Thermo Fisher Scientific). Mobile phase A consisted of water and 0.1% v/v formic acid. Mobile phase B consisted of 80% v/v acetonitrile and 0.1% v/v formic acid. Peptides were loaded at a flow rate of 0.3 μl/min and eluted at 0.2 μl/min or 0.25 μl/min using a linear gradient going from 2% mobile phase B to 40% mobile phase B over 109 or 79 min, followed by a linear increase from 40% to 95% mobile phase B in 11 min. The eluted peptides were directly introduced into the mass spectrometer. MS data were acquired in the data-dependent mode with a 3 s acquisition cycle. Precursor spectra were recorded in the Orbitrap with a resolution of 120,000. The ions with a precursor charge state between 3+ and 8+ were isolated with a window size of 1.6 m/z and fragmented using high-energy collision dissociation (HCD) with a collision energy of 30. The fragmentation spectra were recorded in the Orbitrap with a resolution of 15,000. Dynamic exclusion was enabled with single repeat count and 60 s exclusion duration. The mass spectrometric raw files were processed into peak lists using ProteoWizard (version 3.0.20388) (Kessner et al., 2008), and cross-linked peptides were matched to spectra using Xi software (version 1.7.6.3) (Mendes et al., 2018) (https://github.com/Rappsilber-Laboratory/XiSearch) with in-search assignment of monoisotopic peaks (Lenz et al., 2018). Search parameters were MS accuracy, 3 ppm; MS/MS accuracy, 10ppm; enzyme, trypsin; cross-linker, EDC; max missed cleavages, 4; missing mono-isotopic peaks, 2. For EDC search cross-linker, EDC; fixed modification, carbamidomethylation on cysteine; variable modifications, oxidation on methionine. For sulfo-SDA search: fixed modifications, none; variable modifications, carbamidomethylation on cysteine, oxidation on methionine, SDA-loop SDA cross-link within a peptide that is also cross-linked to a separate peptide. Fragments b and y type ions (HCD) or b, c, y, and z type ions (EThcD) with loss of H_2_O, NH_3_ and CH_3_SOH. 5% on link level False discovery rate (FDR) was estimated based on the number of decoy identification using XiFDR (Fischer and Rappsilber, 2017).

### Integrative structure modelling

#### Input subunits

Using the Mis18α_Yippee_ as a template we generated high-confidence structural models for the Mis18α and Mis18β Yippee domains (using the homology modelling server Phyre2, www.sbg.bio.ic.ac.uk/phyre2/ (Kelley et al., 2015)). These models were almost identical with those obtained using Raptorx (http://raptorx.uchicago.edu/) and AlphaFold2 (Jumper et al., 2021); structure prediction programs that employ deep learning approach independent of co-evolution information (Källberg et al., 2012) (**Fig. 1e**).

#### Scoring function for CLMS

A cross-link was considered satisfied if the Calpha-Calpha distance was less than 22Å. The final score was the fraction of satisfied cross-links.

#### Sampling

To determine the structure of the Mis18 complex we used XlinkAssembler, an algorithm for multi-subunit assembly based on combinatorial docking approach (Schneidman-Duhovny and Wolfson, 2020, Inbar et al., 2005). The input to XlinkAssembler is N subunit structures and a list of cross-links. First, all subunit pairs are docked using cross-links as distance restraints (Schneidman-Duhovny et al., 2005). Pairwise docking generates multiple docked configurations for each pair of subunits that satisfy a large fraction of cross-links (> 70%). Second, the combinatorial assembler hierarchically enumerates pairwise docking configurations to generate larger assemblies that are consistent with the CLMS data.

XlinkAssembler was used with 11 subunits to generate a model for Mis18α/ý: initial hexamer structure based on AlphaFold (Jumper et al., 2021), two Mis18α_Yippee_ domains as well as four copies of the two helices in the Mis18α N-terminal helical region (residues 37-55 and 60-76). For docking Mis18BP1 helices, XlinkAssembler was used with 4 subunits: the Mis18α/ý_Yippee_ domains hetero-dimer and the three Mis18BP1 helices predicted by AlphaFold (residues 21-33, 42-50, and 90-111).

### Cell culture and transfection

The cell line HeLa Kyoto, HeLa 3-8 (having an alphoid^tetO^ array integrated into one of its chromosome arms), as well as HeLa CENP-A-SNAP, GFP Mis18BP1 inducible CENP-A-SNAP and mCherry Mis18α CENP-A-SNAP (kind gift from Iain Cheeseman (McKinley and Cheeseman, 2014)) were maintained in DMEM (Gibco) containing 10% FBS (Biowest) and 1X Penicillin/Streptomycin antibiotic mixture (Gibco). The cells were incubated at 37°C in a CO_2_ incubator in humid condition containing 5% CO_2_. GFP Mis18BP1 was induced with 10 μg/ml doxycycline for 18 h. siRNAs (AllStars Negative Control siRNA 1027280. Mis18α: ID s28851, Mis18β: ID s22367, ThermoFisher) were used in the rescue assays by transfecting the cells using jetPRIME® (Polyplus transfection®) reagent according to manufacturer’s instructions. Briefly, HeLa CENP-A-SNAP, GFP Mis18BP1 inducible CENP-A-SNAP and mCherry Mis18α CENP-A-SNAP cells were seeded in 12-well plates and incubated overnight. siRNAs (50 pmol), vectors (200 ng) and the jetPRIME® reagent were diluted in the jetPRIME® buffer, vortexed and spun down. The transfection mixture was incubated for 15 min before adding to the cells in a drop-by-drop manner. The cells were then incubated for 48 h.

The TetR-eYFP tagged proteins were transfected using the XtremeGene-9 (Roche) transfection reagent according to the manufacturer’s protocol. The HeLa 3-8 cells attached on to the coverslip in a 12-well plate were transfected with the corresponding vectors (500 ng) and the transfection reagent diluted in Opti-MEM (Invitrogen) followed by incubation for 36-48 h.

### Generation of monoclonal antibodies against Mis18α/Mis18

Lou/c rats and C57BL/6J mice were immunized with 60 μg purified recombinant human Mis18α/ý protein complex, 5 nmol CpG (TIB MOLBIOL, Berlin, Germany), and an equal volume of Incomplete Freund’s adjuvant (IFA; Sigma, St. Louis, USA). A boost injection without IFA was given 6 weeks later and three days before fusion of immune spleen cells with P3X63Ag8.653 myeloma cells using standard procedures. Hybridoma supernatants were screened for specific binding to Mis18α/ý protein complex and also for binding to purified GST-Mis18β protein in ELISA assays. Positive supernatants were further validated by Western blot analyses on purified recombinant human Mis18α/ý complex, on cell lysates from *Drosophila* S2 cells overexpressing human Mis18α and on HEK293 cell lysates. Hybridoma cells from selected supernatants were subcloned at least twice by limiting dilution to obtain stable monoclonal cell lines. Experiments in this work were performed with hybridoma supernatants mouse anti-Mis18α (clone 25G8, mouse IgG2b/ƙ) and rat anti-Mis18β (clone 24C8; rat IgG2a/ƙ).

### Western blot

To study the efficiency of DNA and siRNA transfected, HeLa cells were transfected as stated above. Protein was extracted with RIPA buffer and analysed by SDS-PAGE followed by wet transfer using a Mini Trans-Blot® Cell (BioRad). Antibodies used for Western blots were: mouse Mis18α (25G8), rat Mis18β (24C8) (1:100, Helmholtz Zentrum München), Mis18BP1 (1:500, PA5-46777, Thermo Fisher or 1ug/ml, ab89265, Abcam), GFP (1:5000, ab290, Abcam), mCherry (1:1000, ab167453, Abcam) and tubulin (1:2000, T5168, Sigma). Secondary antibodies used were ECL Rabbit IgG, ECL Mouse IgG and ECL Rat IgG (1:5000, NA934, NA931, NA935, GE Healthcare) and immunoblots were imaged using NuGlow ECL (Alpha Diagnostics). For imaging with the Odyssey® CLx system, goat anti-mouse 680 and donkey anti rabbit secondary 800 antibodies were used (1:5000, LI-COR).

### Co-Immunoprecipitation

HeLa Kyoto cells were seeded in 100 mm dishes. The cells were depleted of the endogenous Mis18α or Mis18β by siRNA transfection with jetPRIME® (Polyplus transfection®) and simultaneously rescued with siRNA resistant versions of WT or mutant Mis18α mCherry and Mis18β GFP. The cells were harvested after 48 h and lysed by resuspending in immunoprecipitation buffer, 75 mM HEPES pH 7.5, 1.5mM EGTA, 1.5mM MgCl_2_, 150mM NaCl, 10% glycerol, 0.1 % NP40, 1mM PMSF, 10 mM NaF, 0.3 mM Na-vanadate and cOmplete™ Mini Protease Inhibitor; adapted from (Pan et al., 2017). Cells were incubated with mixing for 30 min at 4°C before sonicating with a Bioruptor® Pico (Diagenode). Lysates were then spun for 10 min at 15,000 *g*. The protein concentrations were determined and adjusted to the same concentration. Protein was taken for inputs, and the rest was incubated with Protein G Mag Sepharose® (GE healthcare), previously coupled to Mis18α antibody, for 1 h at 4°C. Next, the bound fraction was separated from unbound by bind beads to the magnet and washing three times with the IP buffer with either 150mM or 300mM NaCl. The protein was extracted from the beads by boiling with SDS-PAGE Loading dye for 5 min and were analysed by SDS-PAGE followed by Western blotting with anti-mCherry, GFP and tubulin antibodies.

### Immunofluorescence and quantification

The transfected cells were washed with PBS and fixed in 4% paraformaldehyde for 10 min, followed by permeabilisation in PBS with 0.5% Triton™ X-100 (Sigma) for 5 min. The cells were then blocked in 3% BSA containing 0.1% Triton™ X-100 for 1 h at 37°C. The blocked cells were subsequently stained with the indicated primary antibodies for 1 h at 37°C followed by secondary antibody staining under similar conditions. The following primary antibodies were used for immunofluorescence: anti-ACA (1:300; 15-235; Antibodies Inc.) and anti-CENP-A (1:100, MA 1-20832, Thermofisher). The secondary antibodies used were Alexa Fluor® 488 AffiniPure donkey anti-human IgG, Cy5-conjugated AffiniPure donkey anti-human, and TRITC-conjugated AffiniPure donkey anti-mouse (1:300; Jackson Immunoresearch). Vector shield with DAPI (Vector Laboratories) was used for DNA staining.

Micrographs were acquired at the Centre Optical Instrumentation Laboratory on a DeltaVision Elite™ system (Applied Precision) or Nikon Ti2 inverted microscope. Z stacks were obtained at a distance of 0.2 μm and were deconvolved using SoftWoRx, or AutoQuant software, respectively, followed by analysis using ImageJ software. The intensity at the tethering site was obtained using a custom-made plugin. Briefly, the CENP-A signal at the tethering site (eYFP) was found for every z-section within a 7-square pixel box. The mean signal intensity thus obtained was subtracted from the minimum intensities within the section, which was then normalised with the average CENP-A intensities of the endogenous centromeres. The values were obtained from a minimum of three biological repeats. Statistical significance of the difference between normalised intensities at the centromere and tethering region was established by a Mann–Whitney U two tailed test using Prism 9.1.2.

### SNAP-CENP-A assay and quantification

SNAP-CENP-A quench pulse labelling was done as described previously (Jansen et al., 2007). Briefly, the existing CENP-A was quenched by 10 μM SNAP Cell® Block BTP (S9106S, NEB). The cells were treated with 1 μM STLC for 15 h for enriching the mitotic cell population, and the newly formed CENP-A was pulse labelled with 3 μM SNAP-Cell® 647-SiR (S90102S, NEB), 2 h after release from the STLC block (early G1). After pulse labelling, the cells were washed, fixed and processed for immunofluorescence. Images were obtained using DeltaVision Elite™ system (Applied Precision), deconvolved by SoftwoRx and processed by Image J. The average centromere intensities were obtained using a previously described macro CraQ (Bodor et al., 2012). Briefly, the centromeres were defined by a 7×7 pixel box using a reference channel, and the corresponding mean signalling intensity at the data channel was obtained by subtracting the minimum intensities within the selection. The values plotted were obtained from a minimum of three independent experiments. Statistical significance of the difference between normalised intensities at the centromere region was established by a Mann–Whitney U test using Prism 9.1.2.

### Data availability

PDB ID: 7SFY for Mis18α/μ_C-term_ PDB ID: 7SFZ for Mis18α_Yippee_

The MS proteomics data will be deposited to the ProteomeXchange Consortium via the PRIDE (Perez-Riverol et al., 2019) partner repository.

### Code availability

Plugin for analysing intensities at tethering site deposited in Zenodo: DOI 10.5281/zenodo.5708337

## Supplementary Figure Legends

**Supplementary Figure 1** – Mis18α and Mis18β Contain Two Domains Capable of Oligomerising.

**a** & **b**) Domain architecture and amino acid conservation of (**a**) Mis18α and (**b**) Mis18β. Alignments include *Homo sapiens* (*hs*), *Bos taurus* (*bt*), *Mus musculus* (*mm*) and *Gallus gallus* (*gg*). The conservation score is mapped from red to cyan, where red corresponds to highly conserved and cyan to poorly conserved. Secondary structures as annotated/predicted by Conserved Domain Database [CDD] and PsiPred, http://bioinf.cs.ucl.ac.uk/psipred. Multiple sequence alignments were performed with MUSCLE (Madeira et al., 2019) and edited with Aline (Bond and Schüttelkopf, 2009). Dashed boxes highlight Yippee domains whilst solid boxes highlight C-terminus α-helices.

**Supplementary Figure 2** – SAXS Analysis of Mis18α/ý ýN, Mis18α/ý and Mis18_core_ and EDC Crosslinking of Mis18α/ý.

a) SAXS scattering curves of Mis18α/ý ýN, Mis18α/ý and Mis18_core_

b) Guinier Plot showing *Rg* of 53 Å, 60 Å, and 63 Å for Mis18α/ý ýN, Mis18α/ý and Mis18_core_, respectively.

c) Modified Guinier Plot showing *Rc* of 26 Å, 30 Å, and 31 Å for Mis18α/ý ýN, Mis18α/ý and Mis18_core_, respectively.

d) SAXS *P(r)* distributions showing maximum dimensions of 190 Å, 215 Å, and 230 Å for Mis18α/ý ýN, Mis18α/ý and Mis18_core_, respectively.

e) Linkage map showing the sequence position and cross-linked residue pairs between the different Mis18_core_ complex subunits, Mis18α, Mis18β and Mis18BP1_20-130._ Left panel highlights cross-linked residues between Mis18α and Mis18β. Black lines highlight cross-links between N- and C-terminal helical regions of Mis18α. Right panel highlights cross-links observed between i) Mis18BP1_20-130_ and Mis18α (purple) ii) Mis18BP1_20-130_ and Mis18β (light pink) iii) Mis18BP1_20-130_ self cross-links (light grey). White boxes represent residual residues left over from tag cleavage. Dark boxes show Yippee domains and regions of α-helices.

**Supplementary Figure 3** – Structural Characterisation of the Mis18_core_ Complex

**a)** Representative micrograph of negative staining EM of the Mis18α/Mis18β/Mis18BP1_20-130_ (Mis18_core_) complex cross-linked using GraFix (Kastner et al., 2008, Stark, 2010).

**b)** Representative images of 2D classes from Mis18_core_ particles picked using CryoSPARC (Punjani et al., 2017).

**c)** Three models (Class I-III) generated for Mis18_core_ from negative staining EM analysis. All three show that the overall shapes of the Mis18_core_ resemble a telephone handset with ‘ear’ and ‘mouth’ pieces assuming different relative orientations.

**d)** Cartoon representation of the model of Mis18_core_ complex generated in Fig1. Zoomed in panel shows interaction between Mis18α and Mis18β Yippee domains using the second interface. Important residues for this interaction highlighted in pink and purple.

**e)** SEC profile of Mis18α_WT_/Mis18ý_WT_ (red) and Mis18α_C154R/D160R_/Mis18ý_WT_ (black) and corresponding SDS–PAGE analysis of the fractions. Samples were analysed using Superdex 200 increase 10/300 in 20 mM Tris–HCl pH 8.0, 250 mM NaCl and 2 mM DTT.

**Supplementary Figure 4** – Structural and Biochemical Characterisation of Mis18α C-Terminal Helix.

**a)** Cartoon representation of the crystal structure of Mis18α_C-term_/Mis18ý_C-term_ (PDB ID: 7SFY). Mis18α is shown in purple and Mis18β in light pink. Potential residues involved in the interaction are highlighted. Mis18α (purple) and Mis18β (light pink).

**b)** Right panel shows SDS-PAGE analysis of cobalt and amylose pull-down of His-MBP-Mis18β_188–229 WT_ with His-SUMO-Mis18α_191–233_ mutants. SDS-PAGE shows protein bound to nickel resin as input (I) and protein-bound to amylose resin to assess interaction (P). Right panel shows Western blot analysis of co-immunoprecipitation (Co-IP) experiments using Mis18α antibody to test interaction of Mis18α−mCherry with and without mutations in the C-terminal α-helices and Mis18β-GFP. Top panel shows blot against mCherry, middle panel shows blot against GFP and bottom panel shows blot against tubulin as loading control.

**c)** SEC-MALS of His-SUMO-Mis18α_188-233 WT_, His-SUMO-Mis18α_188-233 I201A/L205A_ and His-SUMO-Mis18α_188-233 L212A/L215A/L219A_. Normalised absorption at 280 nm (mAU, left y-axis) and molecular mass (kDa, right y-axis) are plotted against elution volume (ml, x-axis). Measured molecular weight (MW) and the calculated subunit stoichiometry based on the predicted MW. Samples were analysed using a Superdex 75 increase in 50 mM HEPES pH8.0, 150 mM NaCl and 1 mM TCEP.

**d)** Representative immunoblots showing expression levels of endogenous proteins after treatment with siRNA.

**e)** Representative immunoblots showing expression levels of transiently expressed tagged proteins after transfection.

