## Supplementary figures and images for "Structural Basis for Mis18 Complex Assembly: Implications for Centromere Maintenance"

### Figure S1

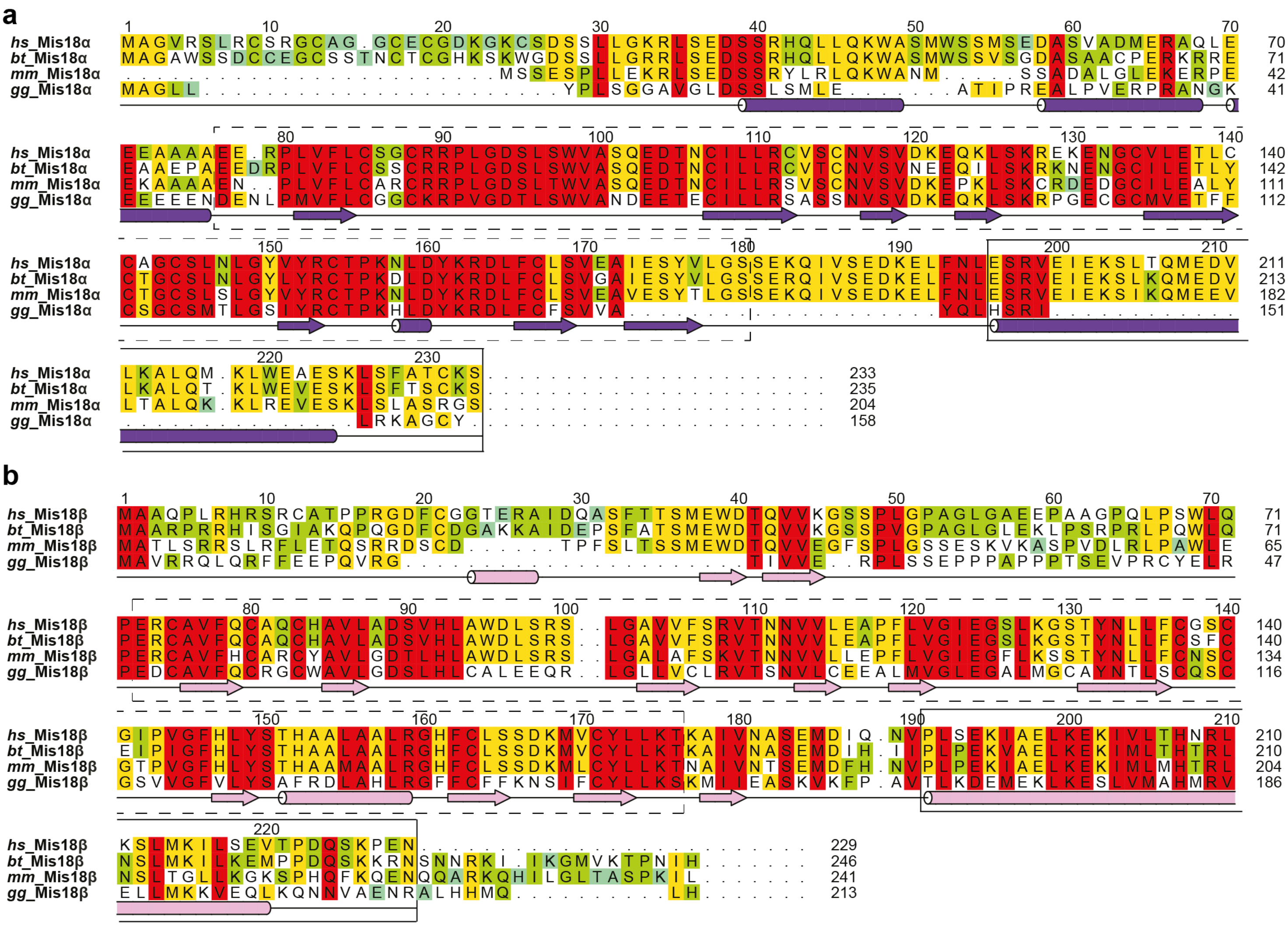

### Figure S2

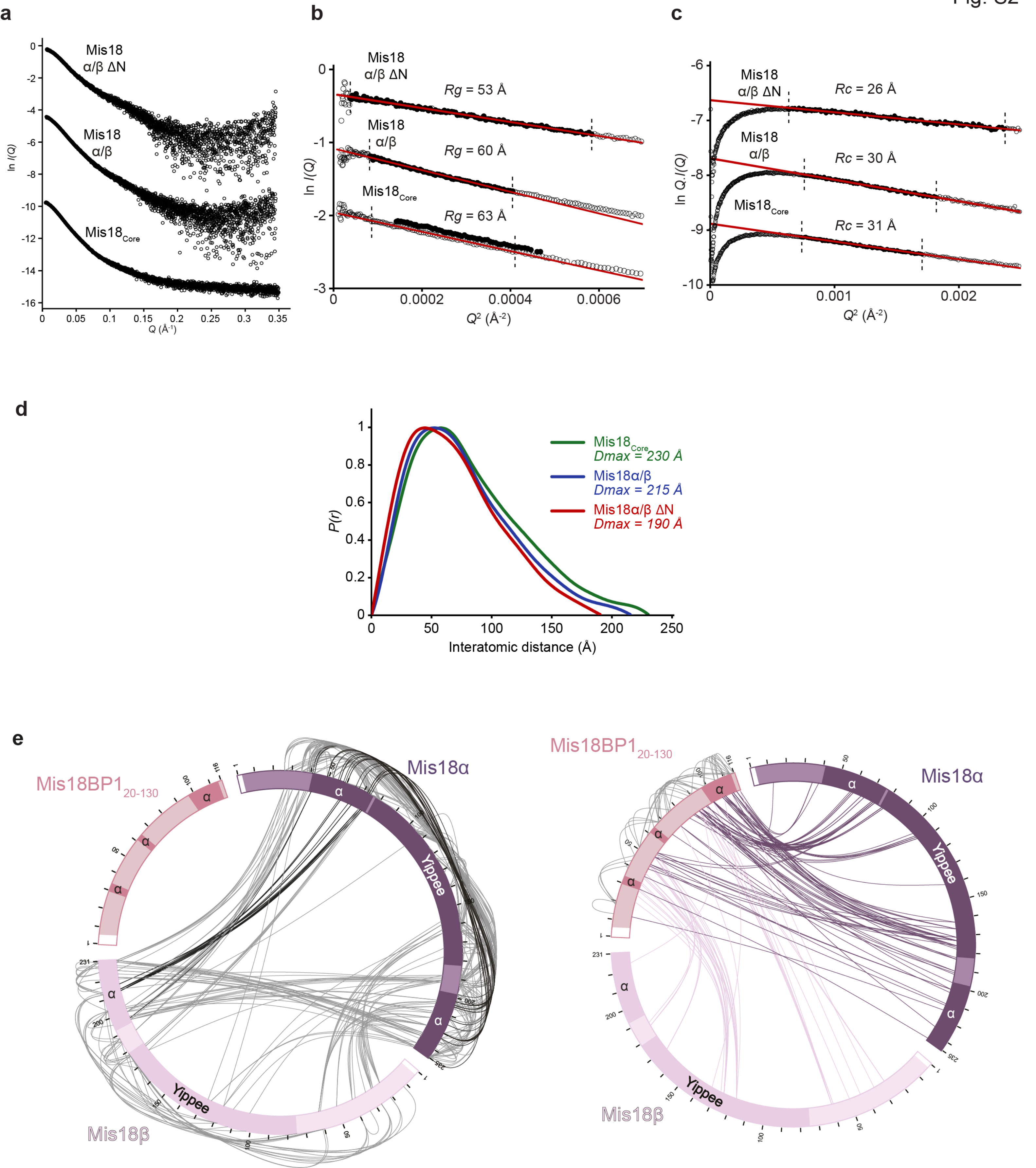

### Figure S3

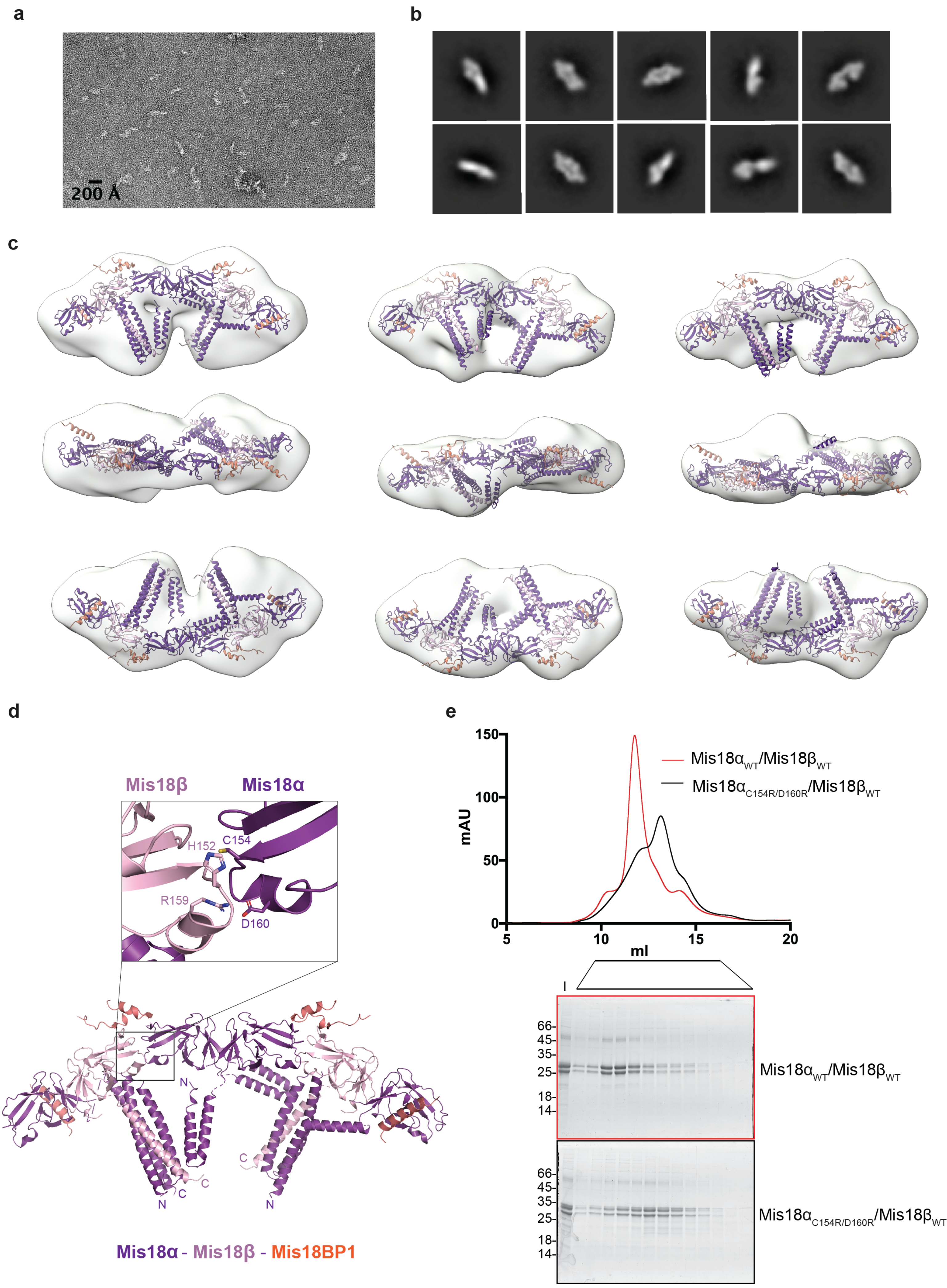
