## Supplementary material for "Structural Basis for Mis18 Complex Assembly: Implications for Centromere Maintenance": Figure S4

Fig. S4

**a**

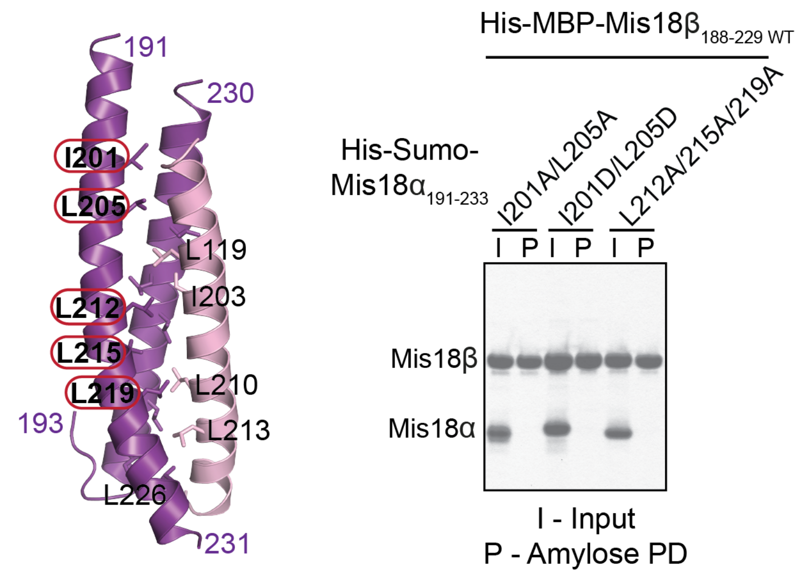


**b**

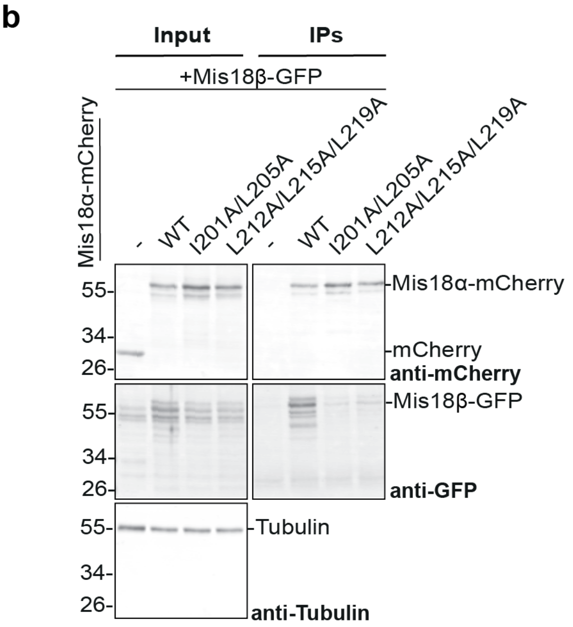


**c**

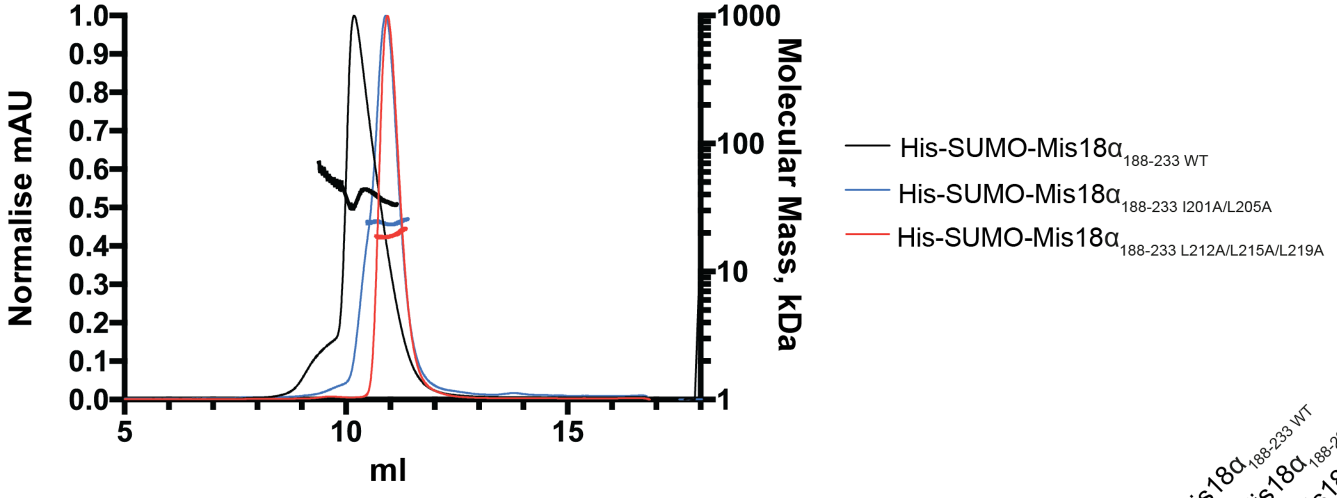


| | Measure MW in kDa (Calculated MW) | Number of Subunits |
|---|---|---|
| His-SUMO-Mis18α$_{188\text{-}233\ WT}$ | 38.8±0.7 (19.5) | Dimer |
| His-SUMO-Mis18α$_{188\text{-}233\ I201A/L205A}$ | 24.1±0.4 (19.5) | Monomer |
| His-SUMO-Mis18α$_{188\text{-}233\ L212A/L215A/L219A}$ | 19.5±0.4 (19.4) | Monomer |

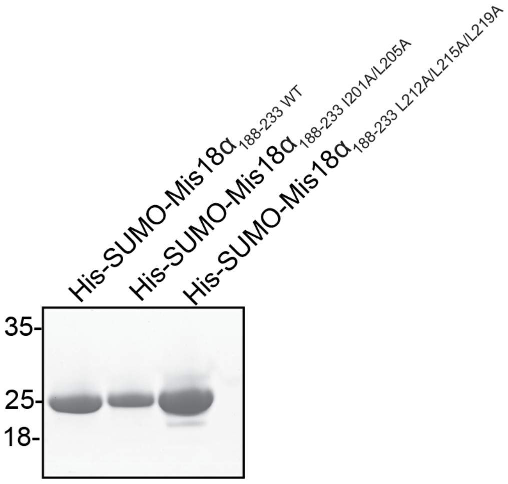


**d**

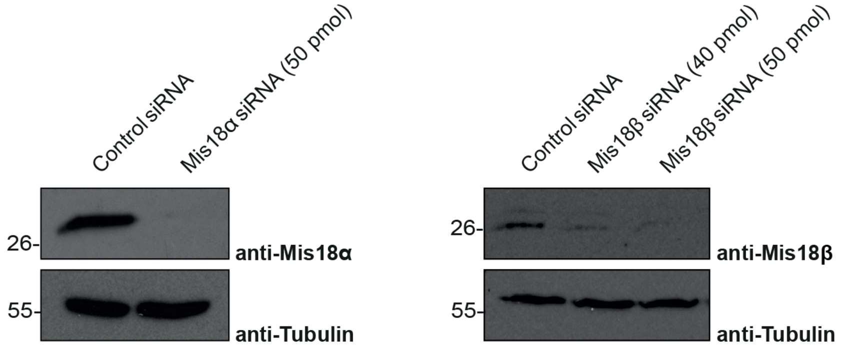


**e**

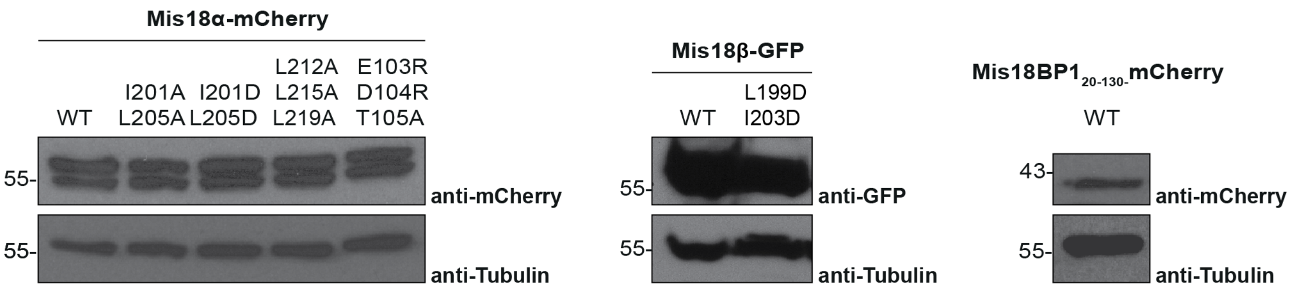
