## Supplementary material for "Structural Basis for Mis18 Complex Assembly: Implications for Centromere Maintenance": Table S1

**Table S1. Crystallographic Data collection and refinement statistics**

| | SeMet-Mis18$\alpha$/$\beta_{C\_term}$ | Mis18$\alpha$/$\beta_{C-term}$ | Mis18$\alpha_{Yippee}$ |
|---|---|---|---|
| Space group | $C222_1$ | $C222_1$ | $P2_12_12$ |
| Unit cell parameters (Å) | a=77.73<br>b=101.91<br>c=88.51<br>$\alpha$=$\beta$=$\gamma$=90° | a=77.591<br>b=101.815<br>c=88.563<br>$\alpha$=$\beta$=$\gamma$=90° | a=110.725<br>b=114.864<br>c=116.279<br>$\alpha$=$\beta$=$\gamma$=90° |
| Wavelength (Å) | 0.97917 | 0.97856 | 1.12713 |
| **<u>Data collection statistics</u>** | | | |
| Resolution range (Å) | 50.00-2.75 (2.75-2.80) | 50.00-2.50 (2.50-2.54) | 50.00-3.00 (3.05-3.00) |
| Number of unique reflections | 9402 (469) | 12467 (600) | 30063 (1479) |
| Completeness (%) | 99.4 (98.3) | 99.0 (96.0) | 99.3 (99.7) |
| $R_{merge}$ | 0.387 (5.327) | 0.078 (0.919) | 0.105 (0.804) |
| $R_{pim}$ | 0.112 (1.456) | 0.040 (0.447) | 0.055 (0.410) |
| Redundancy | 13.4 (12.7) | 5.2 (4.7) | 4.4 (4.5) |
| Mean I/σ | 8.7 (1.5) | 9.2 (1.8) | 16.5 (1.4) |
| **<u>Refinement statistics</u>** | | | |
| Resolution range (Å) | | 44.29-2.50 | 49.99-3.00 |
| $R_{work}$/$R_{free}$ (%) | | 24.77/27.96 | 20.26/25.00 |
| RMSD bonds (Å) | | 0.008 | 0.012 |
| RMSD angles (deg) | | 1.069 | 1.294 |
| Average B factor (Å$^2$) | | 91.18 | 83.84 |
| Number of water molecules | | 17 | 8 |
| Ramachandran favoured (%) | | 97.89 | 95.51 |
| allowed (%) | | 2.11 | 4.49 |
| not allowed (%) | | 0 | 0 |
