## Supplementary material for "Structural Basis for Mis18 Complex Assembly: Implications for Centromere Maintenance": Table S2

**Table S2. Summary of SAXS data**

| | Mis18 α/β ΔN | Mis18α/β | $Mis18_{Core}$ |
|---|---|---|---|
| **SASBDB accession** | | | |
| **Guinier analysis** | | | |
| *I(0)* ($cm^{-1}$) | 0.026 ±7.1x$10^{-5}$ | 0.057 ±1.0x$10^{-4}$ | 0.25 ±6.9x$10^{-4}$ |
| *Rg* (Å) | 53 ±0.21 | 60 ±0.17 | 63 ±0.24 |
| *Rc* (Å) | 26 | 30 | 31 |
| $q_{min}$ (Å$^{-1}$) | 0.0060 | 0.0090 | 0.0093 |
| **P(r) analysis** | | | |
| *I(0)* ($cm^{-1}$) | 0.026 ±6.4x$10^{-5}$ | 0.056 ±1.1x$10^{-4}$ | 0.25 ±6.3x$10^{-4}$ |
| *Rg* (Å) | 55 ±0.17 | 60 ±0.16 | 65 ±0.20 |
| $D_{max}$ (Å) | 190 | 215 | 230 |
| Porod volume (Å$^3$) | 213302 | 377272 | 500371 |
| MW from Porod volume (kDa) | 125 | 222 | 294 |
| $V_C$ (Å$^2$) | 888 | 1161 | 1245 |
| MW from $V_C$ (kDa) | 120 | 192 | 220 |
| ***DAMMIN ab initio* modelling (30 models)** | | | |
| Symmetry | P1 | P1 | P1 |
| *NSD* mean and s.d. | 0.693 ±0.015 | 0.668 ±0.017 | 0.731 ±0.018 |
| $\chi^2$ (reference model) | 1.27 | 1.33 | 0.997 |
| ***DAMMIN ab initio* modelling (30 models)** | | | |
| Symmetry | P2 | P2 | P2 |
| *NSD* mean and s.d. | 0.858 ±0.096 | 0.824 ±0.046 | 0.937 ±0.110 |
| $\chi^2$ (reference model) | 1.26 | 1.33 | 0.976 |
